# Arginine limitation causes a directed DNA sequence evolution response in colorectal cancer cells

**DOI:** 10.1101/2023.01.02.521806

**Authors:** Dennis J. Hsu, Jenny Gao, Norihiro Yamaguchi, Alexandra Pinzaru, Nandan Mandayam, Maria Liberti, Søren Heissel, Hanan Alwaseem, Saeed Tavazoie, Sohail F. Tavazoie

## Abstract

Utilization of specific codons varies significantly across organisms. Cancer represents a model for understanding DNA sequence evolution and could reveal causal factors underlying codon evolution. We found that across human cancer, arginine codons are frequently mutated to other codons. Moreover, arginine restriction—a feature of tumor microenvironments—is sufficient to induce arginine codon-switching mutations in human colon cancer cells. Such DNA codon switching events encode mutant proteins with arginine residue substitutions. Mechanistically, arginine limitation caused rapid reduction of arginine transfer RNAs and the stalling of ribosomes over arginine codons. Such selective pressure against arginine codon translation induced a proteomic shift towards low arginine codon containing genes, including specific amino acid transporters, and caused mutational evolution away from arginine codons—reducing translational bottlenecks that occurred during arginine starvation. Thus, environmental availability of a specific amino acid can influence DNA sequence evolution away from its cognate codons and generate altered proteins.

## Introduction

Genomes of organisms are enriched in certain codons over others. The origins of such codon usage biases have been attributed to both sequence-specific mutational biases that are thought to dominate over long time-scales as well as to organism-specific tRNA availabilities as encoded in the genomes of difference species(*1*). However, the challenge inherent to observing the emergence of such a long time-scale process has precluded definitive support for various proposed models. The mechanisms underlying the emergence of codon usage bias, including the extent to which tRNAs shape genomic evolution have also remained poorly defined. We reasoned that for cancer cells, which divide rapidly, acquire mutations more frequently relative to normal cells, and are exposed to a variety of selective pressures, the evolution of DNA sequence biases would be expedited. This would allow us to detect the emergence of codon-based sequence changes and search for potential underlying mechanisms.

## Results

### Arginine codons and residues are frequently lost through mutation across human cancers

In order to determine if specific codons or amino acids are favored in cancer genomes, we computationally assessed codon-switching events—defined as the gain or loss of a codon via mutation—across all cancers in The Cancer Genome Atlas (TCGA)(*2*). Although dozens of mutational signatures have been shown to be operant with varying weights in different cancers(*3*), we observed that the majority of cancers displayed surprisingly similar patterns of codon gains and losses (**fig. S1A**). Strikingly, when collapsed onto their cognate amino acids, we observed that arginine codons were universally depleted across all cancer types (**Fig. 1A, fig. S1B**). Thus, mutagenic events affecting arginine codons are extremely frequent in cancer.

**Fig. 1.**
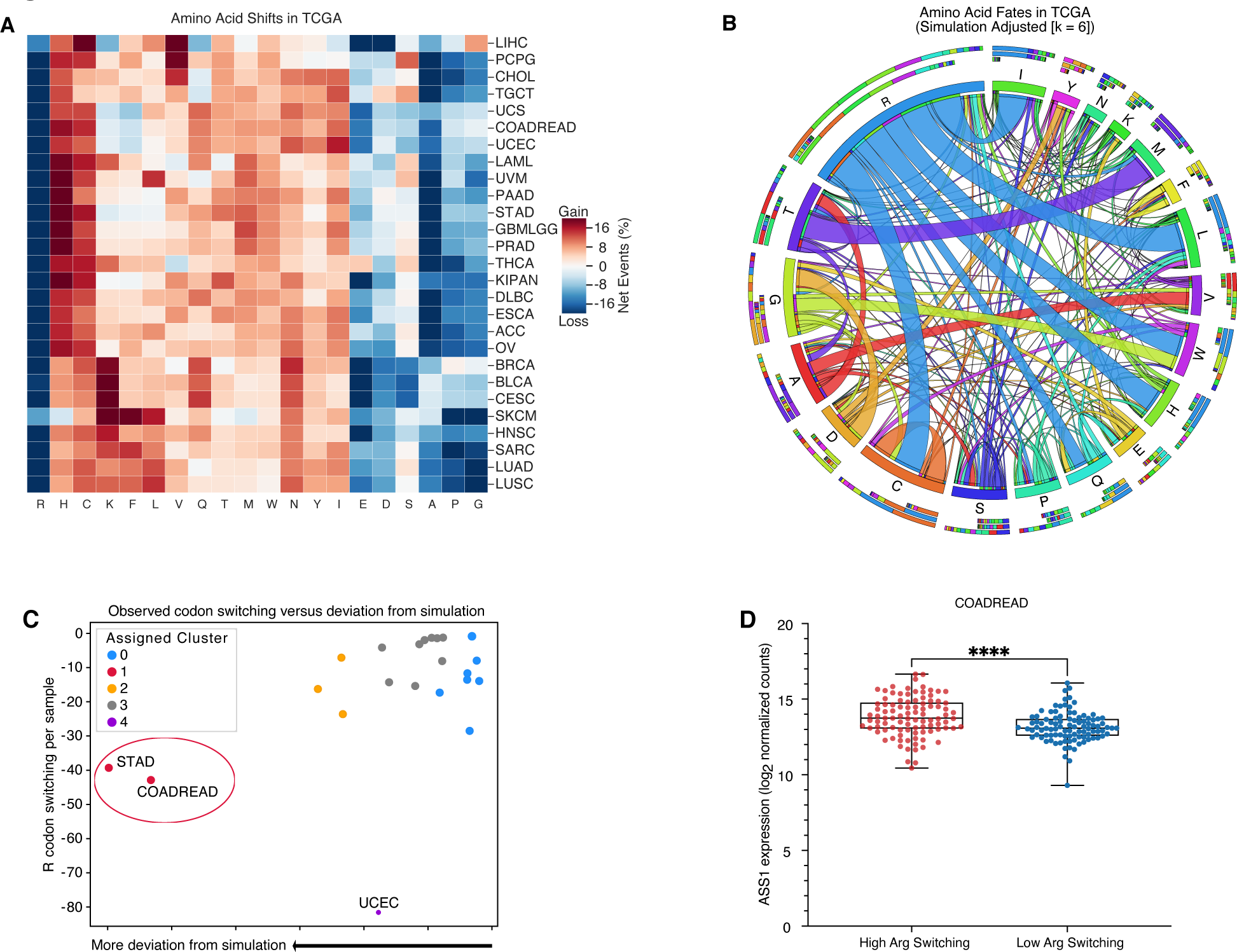
Arginine codons and residues are frequently lost and are associated with an increase in ASS1 expression. **(A)** Heatmap depicting codons gained (red) and lost (blue) across the TCGA. Gains and losses are normalized to the total number of missense and silent mutation events for each cancer type **(B)** Qualitative chord diagram showing amino acid switching events in cancer after adjustment from simulations. Ribbons which directly touch a column segment indicate loss of that specific amino acid codon during a mutational event and gain of the corresponding amino acid codon in which the ribbon terminates. Ribbons that begin and end at the same amino acid represent synonymous mutations **(C)** Arginine codon switching events observed versus predicted. Clusters were assigned with Affinity Propagation **(D)** ASS1 expression in colorectal cancer samples with either a high-degree or low-degree of arginine codon switching events (n=96 per group) (DESeq2 ****P_adjusted_ < 0.0001).

Mutational processes have been shown to act more frequently at specific nucleotides based on their surrounding contexts(*3, 4*). It is therefore important to distinguish between observed codon changes that simply resulted from sequence-specific mutational biases versus codon-switching events that arose from evolutionary selection for or against a given codon. To distinguish between these possibilities, we devised a series of computational simulations that utilized cancer-specific mutational signatures to model expected codon-switching events (**fig. S2**). We extracted mutational signatures from the non-coding regions of cancer genomes to build an unbiased model, reasoning that mutations arising in non-translated regions would be less affected by selective pressures, such as tRNA or amino acid availability, which would be unique to protein-coding genes. Consistent with this possibility, we observed striking disparities in the frequencies of different mutations between coding and noncoding regions of the genome **(fig. S3)**. Modeling codon changes using mutational spectra derived from the non-coding genome further highlighted arginine codon-switching events as being especially overrepresented in coding genes (**Fig. 1B, fig. S4).** These findings support the possibility that certain codon-switching mutational events in the coding genome may confer selective fitness to cells relative to mutations in the noncoding genome. At the codon level, the most frequently lost codons across all cancers were arginine codons: CGG, CGA, CGC, and AGA, with frequent conversions to CAC (histidine), TGC (cysteine), ATA (isoleucine), and CTA (leucine) **(fig. S4)**. Thus, arginine codon-switching events in the coding genome are generally overrepresented even when one considers mutational biases.

Next, in order to identify the tumor types and potential stresses associated with arginine codon mutational loss, we first compared our computational predictions with biological observations for each tumor type. This revealed that stomach adenocarcinoma (STAD) and colorectal adenocarcinoma (COADREAD) tumors were the most enriched in arginine codon-switching events and deviated significantly from the simulated background expectation compared to other cancer types (**Fig. 1C**). In contrast, although endometrial carcinomas (UCEC) exhibited the highest degree of arginine codon-switching, these events were relatively well-accounted for by sequence-specific mutational biases. Thus, the extent to which arginine codon-switching mutations occur in excess vary based on tumor type and are most over-represented in colorectal and stomach cancers.

### Increased arginine codon loss associates with expression of bioenergetic pathways

Because mutations involving arginine codons are especially over-represented in colorectal and gastric cancers, we sought to identify a common pattern between the two. We hypothesized that a potential association with arginine codon loss could be extracellular arginine availability, since arginine is known to become limiting in tumor microenvironments(*5, 6*) and the decoding of arginine codons require this amino acid. Consistent with this, we observed that arginosuccinate synthetase 1 (ASS1), a critical gene that catalyzes the penultimate step of arginine biosynthesis, was overexpressed in both colorectal and gastric adenocarcinoma samples that exhibited high arginine codon-switching events relative to those exhibiting low arginine codon-switching events **(Fig. 1D, fig. S5A)**. Interestingly, ASS1 has been shown to be variably expressed in tumors and can be induced when arginine becomes depleted from the tumor microenvironment(*7, 8*). These data suggest that tumors with increased arginine codon-switching events may have experienced reduced extracellular arginine bioavailability during their development, which would have necessitated the expression of arginine biosynthesis pathway components for survival.

We next asked whether tumors that underwent a high frequency of arginine codon-switching events share common transcriptional programs beyond arginine metabolism. To answer this, we analyzed tumor transcriptomes at a global level using a mutual information-based framework(*9*). We found that in both colorectal and stomach adenocarcinomas, expression levels of genes belonging to S-phase of the cell cycle, DNA replication, nucleotide metabolism, mitochondrial translation, and energetics (glycolysis, the citric acid cycle and electron transport) pathways were significantly correlated with increased arginine codon-switching events **(fig. S6)**. To determine which of these pathways and processes are relevant to the *in vivo* microenvironment, where arginine levels can be significantly limiting(*6, 10, 11*), we conducted a similar analysis on colon and gastric adenocarcinoma cells in the cancer cell line encyclopedia (CCLE)(*12*), where cells were cultured in media containing excess arginine (**Table S1**). Expression of genes belonging to nucleotide metabolism and bioenergetic pathways were selectively modulated in *in vivo* arginine codon-switching tumors but not in cancer cells growing *in vitro* with excess arginine (**fig. S6**). This suggests that provision of arginine to supraphysiologic levels, as is the case for *in vitro* culture of CCLE cells, may reduce cellular dependence on expression of certain bioenergetic (mitochondrial translation, citric acid cycle) and nucleotide metabolism pathways relative to the *in vivo* arginine limiting tumor context.

### Arginine limitation results in nucleotide pool imbalances

Our observations collectively support a model whereby a subset of tumors facing arginine restriction experience perturbations to energy metabolism and nucleotide synthesis. Perturbed nucleotide synthesis can give rise to nucleotide imbalance and in turn increase base misincorporation rates, thereby accelerating mutagenesis and potentiating codon-switching events.

Indeed, arginine metabolism derangements have been shown to impact nucleotide biosynthesis and potentially result in DNA damage(*13–15*). In order to further define the relationship between arginine codon-switching events and arginine and nucleotide metabolism, we collected a panel of colorectal cancer cell lines that were either of the high-arginine codon loss type or the low-arginine codon loss type based on mutational sequence analysis of the CCLE (**Table S2**). We observed that at low concentrations of arginine, within the range reported for tumor core levels(*5*), high-arginine codon loss cell lines exhibited significantly lower viability than low-arginine codon loss lines (**Fig. 2A**). To test whether arginine deprivation results in nucleotide metabolism stress, we performed rescue experiments with extracellular nucleotide supplementation. We observed that while colorectal cancer cell lines exhibited variably impaired growth at low arginine concentrations, there was universal partial rescue of cell viability with nucleotide supplementation **(Fig. 2B, fig. S5B)**. Thus, increased arginine-codon switching is associated with heightened dependence on arginine availability with viability being partially rescued by provision of exogenous nucleotides. Because nucleotide supplementation conferred a survival advantage under low arginine conditions, we sought to define how arginine metabolism affects intracellular nucleotide concentrations. Metabolomic profiling of colorectal cancer cells revealed that arginine deprivation caused depletion of both purine and pyrimidine nucleotides with a greater reduction in high arginine codon mutated lines relative to low arginine codon mutated lines (**Fig. 2C and D, fig. S7)**. It has been suggested that arginine deprivation can impact nucleotide pools through induction of the enzyme asparagine synthetase (ASNS), which converts aspartate to asparagine(*15*). Because aspartate is a critical precursor for nucleotide synthesis, its shunting towards asparagine under arginine starvation would impair nucleotide synthesis. In support of this, arginine deprivation induced ASNS in CRC cells **(fig. S8)**. These findings reveal that arginine deprivation causes nucleotide pool imbalance in CRC cells which contributes to impaired survival.

**Fig. 2.**
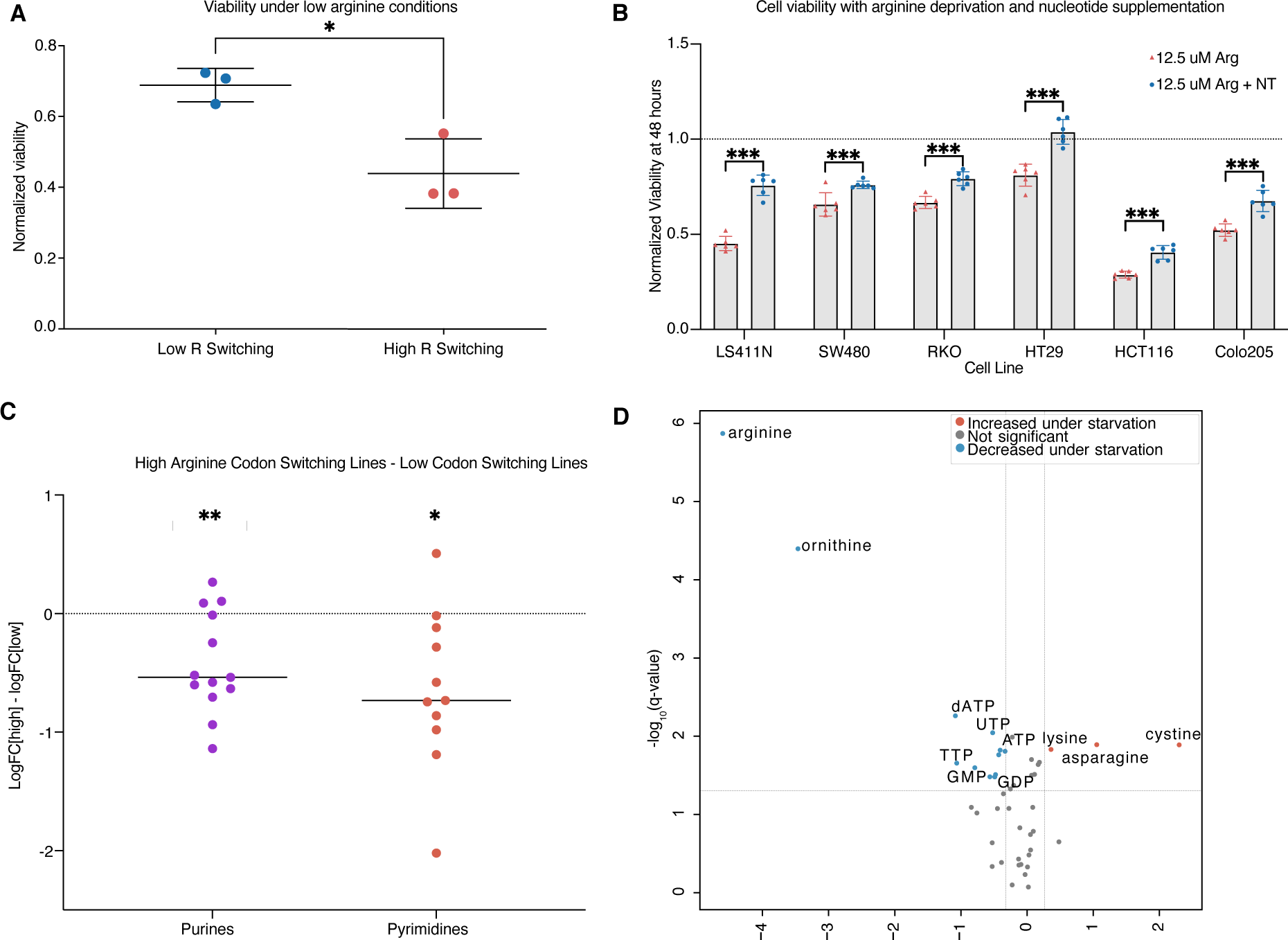
Arginine codon losses are associated with increased dependence on extracellular arginine and nucleotide pool instability during starvation. **(A)** Cell line viability under low arginine (12.5 uM) conditions. (n = 3 per group) **(B)** Effect of nucleotide supplementation on colon cancer cell viability with arginine deprivation (n = 6 per group, two-tailed t-test) **(C)** Metabolite profiling differences after exposure to low arginine concentrations for 24 hours. Each point represents a purine/pyrimidine pathway metabolite and is the average log2FC difference between high arginine codon-mutated lines and low-codon mutated lines (one-sample t-test with μ0 = 0). **(D)** Volcano plot of metabolite changes following arginine deprivation. Only detected citric acid cycle, urea cycle, amino acids, and nucleotide intermediates are labeled. (*P < 0.05, **P < 0.01, ***P<0.001)

### Arginine limitation causes an acute arginyl tRNA repression response

Our findings reveal that arginine limitation of CRC cells that are more reliant on extracellular arginine alters nucleotide pool balance, which can potentially cause mutations. These findings, however, do not explain why arginine codon-switching events are enriched in specific tumors. We thus focused on the association between arginine availability and arginine codon-switching events. Metabolic perturbations such as oxidative stress and glutamine deprivation were recently shown to reduce the levels of specific charged tRNAs—inhibiting translation of downstream genes(*16, 17*). Furthermore, complete elimination of arginine from the environment has been shown to induce ribosome pausing at arginine codons in bacteria as well as in mammalian cells *in vitro* – repressing global protein synthesis(*18, 19*). Because arginine codon-switching mutations would theoretically lessen the requirement for arginine tRNAs during protein translation, we hypothesized that arginine codon switching events may facilitate gene expression under conditions where arginine becomes limiting. Indeed, arginine codon-switching mutations tended to occur in higher-expressed genes in patient samples, highlighting the possibility that arginine codon-switching events might have an outsized influence on gene translation **(fig. S9)**. In such tumors, switching to non-arginine codons may facilitate gene translation in contexts where environmental arginine becomes limiting. We therefore sought to quantify how arginine deprivation affects availability of arginine tRNAs. We assessed tRNA levels in colorectal and gastric cancer cells following arginine starvation through northern blotting. Remarkably, arginine limitation acutely and dramatically depleted arginine tRNA levels (**Fig. 3A**). We detected a significant reduction in multiple arginine tRNA isodecoders including tRNA^Arg^_UCG_, tRNA^Arg^_UCU_, and tRNA^Arg^_CCG_ within 24 hours of arginine deprivation. Importantly, we did not observe reduced levels of other tRNAs such as tRNA^Leu^_UAG_, tRNA^Tyr^_GUA_, or tRNA^His^_GTG_ upon arginine restriction. Thus, extracellular arginine restriction causes an acute and substantial reduction of arginine tRNA levels in CRC cells.

**Fig. 3.**
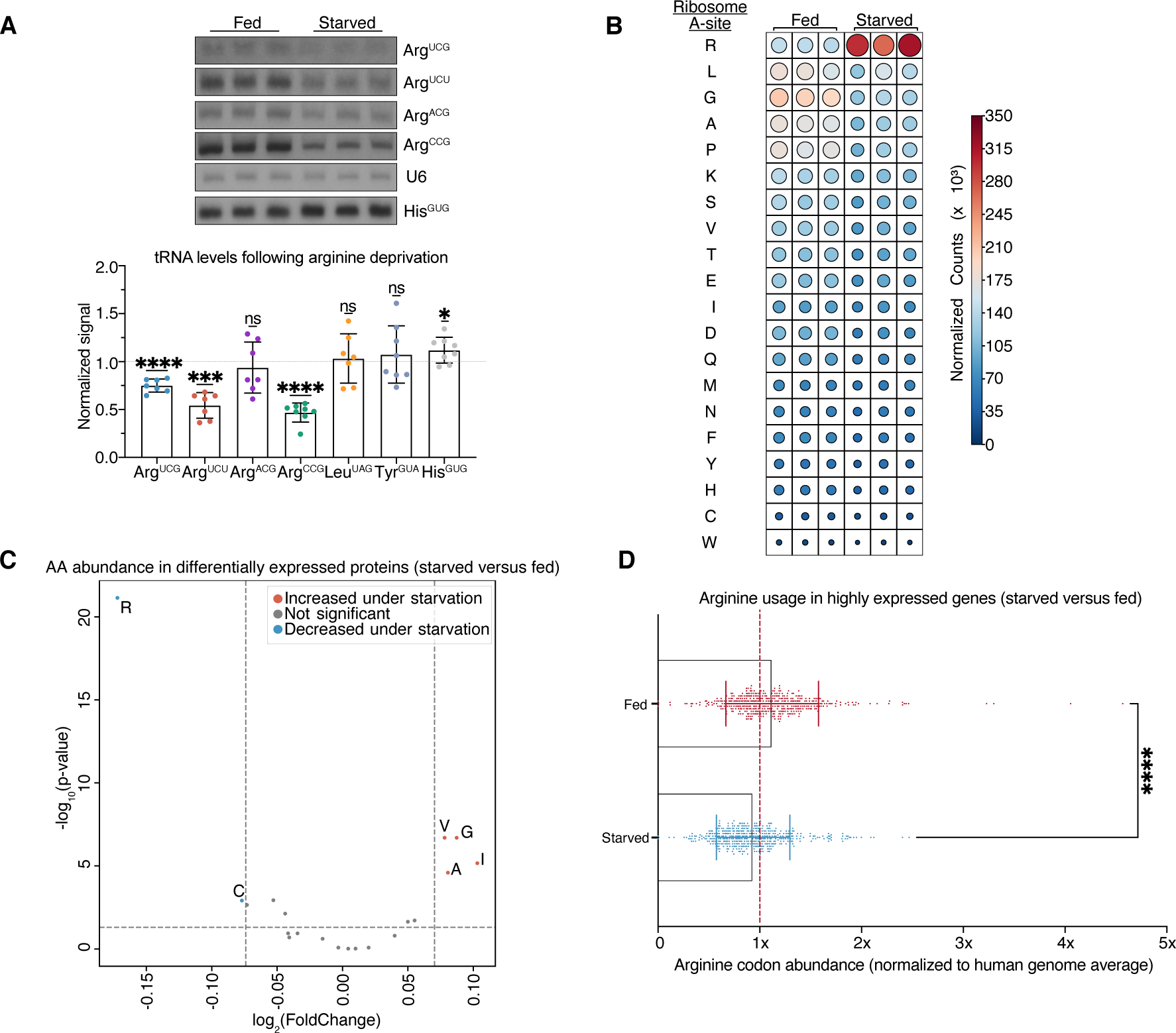
Arginine deprivation reduces arginine tRNA availability, increases arginine ribosome localization, and reduces arginine usage in the tumor proteome. **(A)** tRNA quantification as assessed via northern blot. Each dot represents the average abundance in an independent colon cancer or gastric cancer cell line (n = 7 per group, one-sample t-test with μ0 = 0) **(B)** Ribosome A-site localization counts from ribosomal profiling experiments under starved or fed conditions. Circle size is scaled to counts **(C)** Amino acid usage in genes that are highly expressed under fed or starved states **(D)** Arginine codon abundance in genes expressed in fed or starved states (n ≥ 450 per group, two-tailed Mann-Whitney test). Proteins are stratified based on the top 10% most changed in either fed or starved states. (*P<0.05, **P<0.01, ***P<0.001, ****P<0.0001)

We hypothesized that the repression of arginine tRNAs may be caused by reduced tRNA aminoacylation in the setting of arginine limitation. Reduced aminoacylation of certain tRNAs has been shown to destabilize tRNAs(*20*). To test this, we inhibited aminoacylation in CRC cells by depleting the arginyl-tRNA synthetase RARS and quantified tRNA levels (**fig. S10**). Indeed, suppressing arginine aminoacylation significantly suppressed expression of multiple arginyl tRNAs. These findings are consistent with arginine limitation causing reduced arginyl-tRNA charging and consequently, degradation or destabilization of arginyl tRNAs.

### Arginine restriction causes ribosomal stalling at specific arginine codons

Significant reductions in arginine tRNA availability would be expected to impair arginine-codon dependent translation. To quantify how arginine deprivation-mediated tRNA changes impact gene translation, we performed ribosomal profiling (*21, 22*). As expected, arginine starvation significantly increased ribosomal occupancy at arginine codons under starvation conditions (**Fig. 3B**). Such increased ribosomal A-site localization over arginine codons upon arginine restriction is consistent with increased stalling at arginine codons. As orthogonal approaches for assessing ribosomal dynamics, we utilized two additional metrics to quantify ribosome stalling events. We first calculated Consistent Excess of Loess Predictions (CELP) coefficients to measure the degree of stalling at all codons(*23*). This analysis further confirmed global and significant increases in stalling at arginine codons upon arginine deprivation **(fig. S11A).** Second, we calculated the frequency of amino acid appearances immediately upstream and downstream of maximal ribosome stalling sites during arginine deprivation and observed that arginine codons were significantly over-represented near the global stalling maxima of transcripts, on average appearing more than twice as often as expected **(fig. S11B)**. In contrast, we found no such evidence of arginine enrichment near stalling sites during arginine replete conditions **(fig. S11C).** These findings reveal that arginine deprivation significantly increases ribosome stalling events at arginine codons.

Next, in order to understand how codon-switching events influence ribosome dynamics, we compared ribosome localization in specific genes that were heterozygous for single nucleotide variants (due to arginine codon-switching at one allele) under arginine-fed and arginine-deplete conditions. This experimental model provided us wild-type and mutant arginine codon endogenous ‘reporters’ for specific genes. Remarkably, variant alleles that underwent codon-switching away from arginine codon usage showed less ribosome stalling under arginine limitation at those specific codon positions compared to their corresponding wild-type alleles in the same cell **(fig. S12A, fig. S13).** By comparison, codon-switching events that only involved non-arginine codons showed no significant differences in ribosome stalling at the wild-type versus variant alleles **(fig. S12B)**. Consistent with these observations, genes that harbored arginine codon changes generally showed less stalling at multiple arginine codons under arginine starvation conditions **(fig. S14A)** and consequently higher translational efficiency compared to genes that were wildtype with respect to arginine codon mutation status **(fig. S14B).** These results demonstrate that codon-switching events—specifically, the loss of rate-limiting codons—can directly influence ribosome localization dynamics under amino acid limitation. Therefore, whereas a direct consequence of arginine starvation-mediated tRNA changes is increased ribosome stalling at arginine codons, mutations that result in a loss of an arginine codon tend to relieve this translational bottleneck.

### Arginine limitation causes a proteomic shift from arginine rich to arginine low proteins

Significant stalling of arginine translation would be predicted to alter arginine utilization in the tumor proteome. We thus performed TMT-based quantitative proteomics under conditions of arginine excess versus limitation and found that arginine deprivation resulted in a shift in the tumor proteome towards proteins with substantially lower arginine content **(Fig. 3C)**. Moreover, arginine usage in highly expressed genes was highly significantly reduced upon arginine restriction **(Fig. 3D)**. Pathway enrichment analysis of proteomic changes revealed that proteins related to amino acid transport and DNA damage-induced senescence were increased, whereas proteins related to DNA strand elongation and interferon alpha/beta signaling were reduced upon arginine restriction **(fig. S15A).** Notably, proteins that were upregulated in these pathways upon starvation also tended to show reduced arginine codon content (**fig. S15B**). At the codon level, the three most affected codons with respect to under-utilization in proteins were all arginine codons **(fig. S16)**. Thus, arginine deprivation promotes induction of multiple gene expression programs that utilize arginine less frequently. The significantly decreased need for arginine within multiple gene sets responding to arginine limitation, such as amino acid transport and synthesis, suggests that the amino acid requirements for expression of these gene sets may have undergone prior selection to allow the continued expression of specific stress response programs when arginine is limiting. When coupled with the ribosomal profiling data, these findings suggest that during arginine scarcity, when arginine tRNAs become limiting, there may be an evolutionary advantage for tumors that have undergone additional codon-switching events from arginine codons to codons for which cognate tRNAs remain available for usage in translation.

Our findings thus far reveal that arginine restriction causes an acute response whereby arginyl tRNAs become repressed, leading to ribosomal stalling at rate-limiting arginyl codons of highly expressed genes. This is associated with a proteomic shift away from arginine-rich proteins towards arginine-low proteins, which includes amino acid transporters. Arginine restriction also causes nucleotide imbalance, accelerating mutagenesis. We hypothesized that over longer timescales, this context selects for cancer cells that have undergone arginine codon mutational switching events, which enables translation of proteins that are adaptive for survival.

### Arginine restriction is sufficient to cause arginine codon-switching evolution *in vitro*

We next sought to determine if arginine restriction is sufficient to causally drive codon-switching events away from arginine. To do this, we conducted laboratory evolution experiments by culturing colon cancer cells under reduced arginine conditions and assessed genomic codon-switching events using whole exome sequencing (**Fig. 4A**). Iterative passaging of multiple colorectal cancer cell lines over eight passages (∼24 population doublings lasting ∼2 – 3 months) caused a significant increase in arginine codon-switching events in arginine restricted cells relative to control cells that were passaged the same number of times under arginine-rich conditions **(Fig. 4B)**. Consistent with our prior observations that arginine deprivation results in nucleotide pool imbalances, potentially accelerating mutational rate, arginine deprivation was associated with an increase in general mutational load **(fig. S17)**. Strikingly, arginine codon mutations were increased in genes upregulated during arginine deprivation, as identified from our prior proteomics experiments, compared to genes that were highly expressed during the arginine fed state **(Fig. 4C)**. In contrast, the rate of histidine mutational losses between the gene sets were not significantly different **(Fig. 4D)**. Thus, arginine deprivation *in vitro* is sufficient to increase the frequency of arginine codon-switching mutational events.

**Fig. 4.**
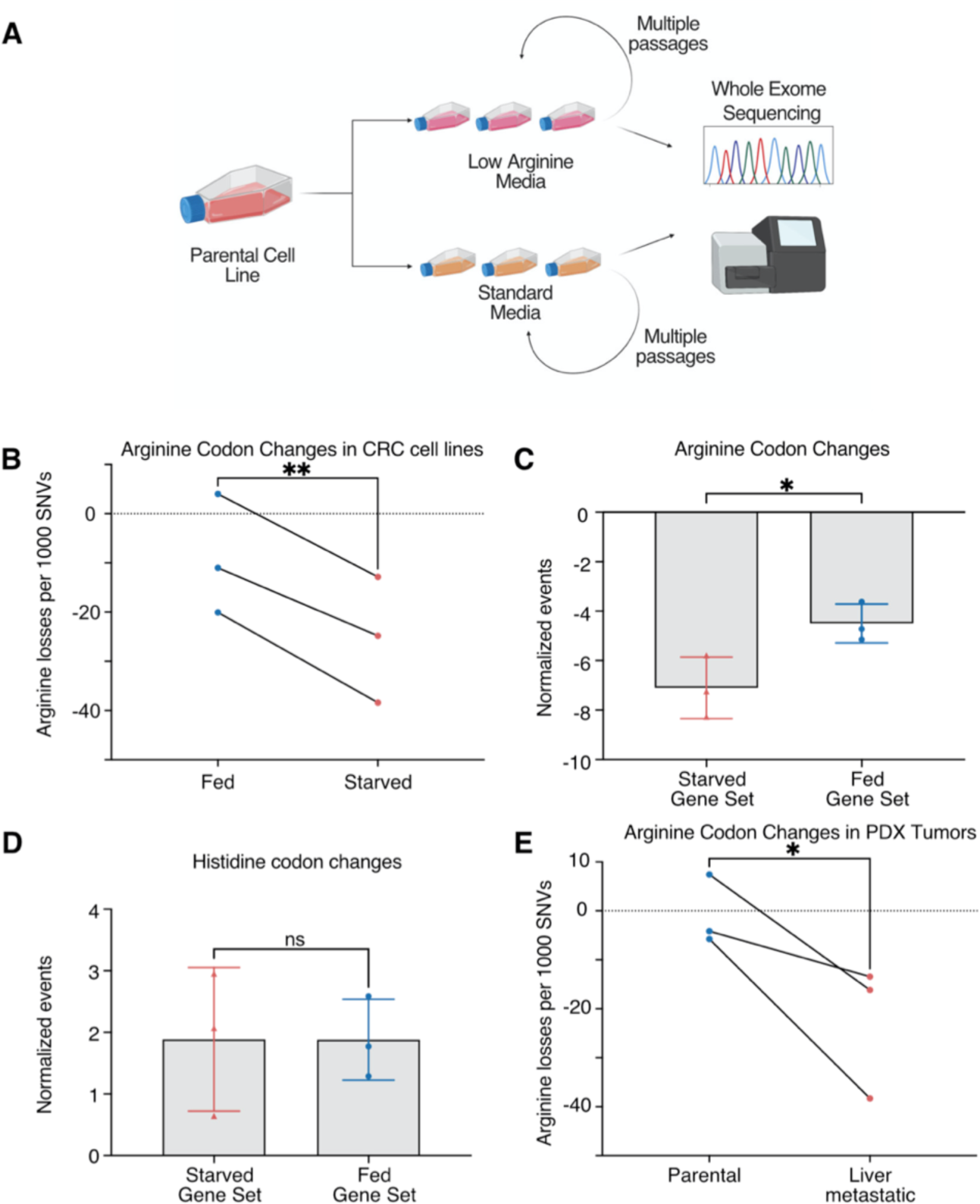
Arginine deprivation promotes arginine-losing mutations. **(A)** Schematic of arginine deprivation experiments. **(B)** Arginine codon changes in cells serially passaged in either full media or low arginine media (n=3 per group, two-tailed paired t-test). **(C)** Arginine codon changes in proteins that are increased during the fed or arginine-starved states (n = 3 per group, two-tailed t-test). **(D)** Histidine codon changes in proteins that are increased during the fed or arginine-starved states (n = 3 per group, two-tailed t-test) **(E)** Arginine codon changes in PDX tumors that underwent multiple rounds of in vivo liver metastatic selection (n = 3 per group, one-tailed paired t-test). (*P<0.05, **P<0.01)

### Arginine limitation causes arginine codon-switching evolution *in vivo*

We next asked if we could recapitulate arginine codon-switching events *in vivo* and whether tumor propagation in a microenvironment low in arginine could also elicit such codon-switching events. We specifically focused on the liver microenvironment due to the liver being the organ in which the arginine degrading enzyme, arginase, is most highly expressed(*24*), and also because the liver is a frequent and pathophysiologically relevant site of distant organ metastatic relapse in both colorectal and gastric cancers(*25, 26*). We first analyzed metabolite profiling data of highly liver metastatic patient-derived xenograft tumors that had undergone at least 5 rounds of *in vivo* selection for liver colonization(*27*) and observed that arginine was indeed the lowest abundance free amino acid in the highly liver metastatic tumors compared to the parental tumors (**fig. S18 A and B)**. We next conducted whole exome sequencing of additional PDX tumors and observed that the rate of acquisition of new arginine codon mutations was significantly increased in tumors that had undergone serial rounds of *in vivo* liver colonization selection compared to the rate measured in the parental tumors **(Fig. 4E)**. Thus, reduced arginine bioavailability *in vivo* is associated with an increased rate of arginine codon mutations, mirroring our observations *in vitro*. These findings as a whole reveal that limitation of a single amino acid, arginine, results in multiple consequences in colorectal cancer cells. First, arginine limitation causes nucleotide pool imbalances and increased mutational rate. Concurrently, arginine tRNA levels are reduced, resulting in ribosomal stalling at arginine codons, providing a selective pressure against arginine codon usage and providing an evolutionary advantage to cancer cells whose coding genomes require less arginine for translation of highly expressed genes required for growth. This context selects for cancer cells that have undergone arginine codon mutational switching events in the coding regions of such growth promoting genes. Based on the totality of these observations, we propose that limitation of an amino acid (arginine) can causally increase the rate of mutations of its cognate codons in the cancer genome—facilitating the continued translation of proteins that may be adaptive for responding to the specific amino acid restriction and leading to the generation of novel proteins with arginine substitutions.

## Discussion

The acquisition of somatic mutations contributes to the development of cancers *de novo,* the emergence of treatment resistance, and can predict response to immunotherapy(*28–31*). Understanding the mechanisms that drive new mutations remains an important problem in cancer biology and oncology. Environmental contributions to mutational processes have generally been thought of as foreign additions to a system: examples include UV radiation, tobacco smoke, or aristolochic acid. Our work suggests that environmental deprivation, i.e., absence or restriction, of just a single amino acid can drive switching away from specific codons in the human cancer genome by simultaneously enhancing mutagenesis and altering specific cognate tRNA availability. In yeast, genetic defects in nitrogen metabolism can increase mutational rates in strains with heightened mutagenic backgrounds(*32*). Moreover, genetic defects in the urea cycle, a critical downstream pathway in the utilization of intracellular arginine, can result in altered rates of pyrimidine synthesis and affect mutational spectra(*13, 14*). While these studies have focused on genetically driven defects in metabolism resulting in mutagenesis, our findings reveal that availability of a specific environmental nutrient, arginine, can filter the mutational landscape of cancer cells in a codon-dependent manner and drive them towards acquisition of arginine-codon mutations. Others have also shown that for a given tRNA, distinct isodecoders associate with proliferation versus differentiation states(*33*). Our findings of a rapid depletion of arginine tRNAs upon arginine limitation as well as the induction of an arginine-low tumor proteome suggest the existence of an acute tRNA-mediated stress response to arginine restriction that promotes translation of genes with reduced arginine codons. We find that in colon cancer cells, limitation of arginine causes an acute translational shift towards an arginine low proteome. Prior work in bacteria and in mammalian cells *in vitro* had shown that complete elimination of arginine from the environment can cause ribosome pausing and global translational repression that was proposed to be caused by reduced tRNA aminoacylation(*18, 19*). Our findings across a series of colon cancer cell lines reveals that physiological limitation of arginine represses the levels of arginine tRNAs, an effect that could also be elicited upon repression of arginyl tRNA aminoacylation. Our findings are thus consistent with a combination of arginyl tRNA repression and reduced aminoacylation contributing to ribosomal pausing at cognate arginine codons and inducing a proteomic shift in response to arginine deprivation in colon cancer cells.

To our knowledge, this is the first demonstration of directed DNA evolution and selection against specific codons in response to a specific environmental perturbation. Our observations imply that over time, cancer cells growing in an arginine-scarce environment are likely to lose more arginine codons and suggest that *in vitro* systems currently used to study cancer and other diseases, for example cells growing in tissue culture, are potentially susceptible to evolving away from arginine codons at different rates depending on their level of arginine supplementation, the fidelity of their DNA repair mechanisms, and the robustness of their arginine tRNA pool. Further work is required to understand if other nutrient limitations also elicit DNA sequence evolution and to understand how other genetic and environmental factors, such as competition with the surrounding microbiome or presence of inflammatory states interact with nutritional availability to affect DNA sequence evolution. With respect to arginine, it has already been suggested that free arginine is especially critical for regulating cancer immunology(*34*), thus competition for this common substrate may influence the evolution of the cancer genome, especially in contexts of tumors with high immune infiltration. Inflammatory bowel disease and H. Pylori infections, precursor disease states with established epidemiological and pathophysiological links to the development of colorectal and gastric cancer, respectively, have both been shown to modulate arginine availability in affected tissues(*35, 36*). In fact, recent work has elegantly demonstrated that amino acid limitation can be so significant under inflammatory-signaling that cancer cells utilize alternative translational decoding for specific amino acids(*37*), leading to the production of altered proteins and neo-antigens. Our findings reveal that limitation of an amino acid can also elicit protein sequence changes and perhaps neo-antigen load via an alternative mechanism—DNA sequence codon-switching events. Notably, significant arginine limitation superimposed on a background of increased base misincorporation rates, such as in mismatch repair deficiency would increase the probability of stochastically acquiring arginine codon mutations which may then confer a survival advantage, and may partially contribute to the increased signal in some tumors over others. However, our experiments reveal that this process occurs in both mismatch repair proficient and mismatch repair deficient tumors and cell lines. Finally, our findings reveal that codon-based mutations can potentially identify subsets of cancers that are more sensitive to restrictions of a specific amino acid. These findings have implications for dietary amino acid restriction approaches that have been tested in tumor models as well probiotic engineering approaches that can modulate tumoral amino acids(*38–42*). The codon-based genotype-dependent vulnerability described herein suggest potential for use of codon-centric mutational spectra as biomarkers for emerging cancer metabolism oncologic therapies.

## Materials and Methods

### Experimental design

Sample sizes were selected back on knowledge of intra-group variation and expected effect size. For *in vitro* experiments, sample sizes were chosen based on prior knowledge on intra-group variation. Data was collected based on pre-determined end-points (in vitro assays) or tumor burden exceeding 2000 mm^3^. Experiments were carried out in replicates as noted in the text and figure legends. Samples were allocated randomly if possible. No blinding was performed.

### Codon mutation analyses

Mutation annotation files (.maf) corresponding to TCGA studies were downloaded from the Broad Firehose platform (http://firebrowse.org). When possible, we utilized combined cancer data sets (i.e. COADREAD, KIPAN, and GBMLGG). A script was written in Python (V 3.8.5) to manually count codons lost and gained across the coding regions of cancer-types and samples. For each cancer sample, the count for a codon was subtracted if it was lost through a missense or silent mutation and added if it were gained through a silent mutation. An “event” was defined as the gain and loss of a pair of codons. We also utilized a similar framework to count the total flux between codons or amino acids to determine flux between codons and amino acids. Events were plotted using circos plots(*43*).

#### Derivation of null distributions

We utilized a Monte Carlo approach to derive various null distributions of codon and amino acid usage shifts across cancer types and different samples. Briefly, our algorithm scatters nucleotide mutations across reference gene sequences downloaded from ENSEMBL. Prior to input into the simulation, genes with multiple splice variants were filtered against the APPRIS database to include only the highest-ranking principle splice variant for simulation(*44*). Probabilities of specific nucleotide mutations were weighted based on the 5’ and 3’ contexts of each nucleotide(*3*).

For inputs into the analysis, we first downloaded the mutation calls from ICGC (last accessed February 2018) (*45*) and then cross-referenced the intergenic and intronic mutational calls with the reference genome to extract the 5’ and 3’ nucleotide contexts to infer mutational probabilities of different nucleotides under different contexts. Each cancer type was assigned its own unique mutational matrix. For each tumor sample in the TCGA, we created a corresponding *in silico* sample and constrained potential mutations to the same set of genes that are mutated in each specific TCGA sample. For each *in silico* sample, candidate genes were randomly mutated the same number of times as was observed in its matched TCGA sample. These specific constraints were placed to prevent the model from deviating due mutations being simulated on lowly mutated genes or genes with wildly different codon content compared to the original sample. For each gene, nucleotides positions are first hashed by 5’- and 3’- contexts and selected for mutation using a vectorized approach to randomly select possibilities along the entire transcript. The effect on the gene (codon change and amino acid change) were calculated and utilized for downstream analyses. Each sample was simulated a thousand times (n=1000).

For statistical inference, we created a “null distribution” mean for different codon/amino acid gains/losses by populating the dataset with mean inferences from each individual TCGA sample. A log-rank test was then performed to determine the extent to which the observed TCGA dataset was different from the simulated dataset. Heatmaps were generated using the Seaborn library in Python (seaborn). Chord diagrams were generated using both the observed datasets and simulation data using Circos(*43*). Qualitative circos plots were generated using scaled values following the formula provided by the developer using the following equation: (e^k*x/max(x)^ – 1) / (e^k^ – 1), where *k* is the scaling factor, x is ratio of the observed shift to the simulation mean for the specific shift, and max(x) is the maximum test-statistic across the entire simulation.

### Gene expression analysis in arginine-codon switching samples

Tumors from TCGA were assigned as high or low arginine codon switching using the *in silico* model described above and assigned a z-score based on the number of deviations from expectation for each sample. The top and bottom 20% of samples were assigned as high-switching and low-switching respectively. To determine if ASS1 expression is differentially expressed between tumors with high or low arginine codon-switching, raw counts from RNA sequencing were obtained using the TCGAbiolinks package in R(*46–48*) and subsequent normalization and differential gene expression analysis between high and low arginine codon-switching groups performed using DESeq2(*49, 50*). For graphical purposes, DESEq2 log-normalized counts are plotted with the DESeq2 p-value (adjusted for multiple comparisons across all genes) annotated for ASS1.

In order to contrast gene expression patterns between *in vivo* and *in vitro* cancer cells, spearman correlation coefficients were calculated between median-of-ratios normalized count data and the codon-switching scores assigned from simulation, then ranked based on strength of correlation for mutual information analysis with iPAGE(*9*). For CCLE samples, RNA sequencing count data were obtained from the Cancer Dependency Map version (most recently processed with release 21Q4) (*51, 52*) and also normalized with median of ratios using DESeq2. Codon changes in colorectal and gastric cancer cell lines were calculated based on corresponding mutational data which were obtained from the Cancer Dependency Map. Spearman correlation coefficients between gene expression and arginine codon loss were calculated and then input for mutual information analysis identical to how the TCGA samples were processed. For the iPAGE program, the independence flag was set to zero to allow for calculation of over-representation in the maximum number of pathways and the ebins parameter set to four. To graphically depict shared pathways, only genes in the top bin (corresponding to the top 25% of correlated genes) with pathway over-representation in both COADREAD and STAD datasets were selected for graphing, with pathways collapsed onto the most top-level statistically significant pathway in the Reactome hierarchy(*53*).

#### Analysis of mutational events in TCGA RNAseq data

Raw counts from TCGA were obtained using TCGAbiolinks package in R(*46–48*). Gene size was estimated using the GenomicFeatures package in R (*54*) to calculate gene size from exon length using the GRCh38.105.gene transfer format data from ENSEMBL and used the calculate transcripts per million for each gene in each sample. Genes were then sorted and ranked within each sample. Mutational events in the top or bottom of half of gene expression in each sample were counted by cross-referencing and matching sample barcodes to whole-exome sequencing data collected in the TCGA. High-arginine mutated and low-arginine groups were assigned as previously specified.

#### Arginine viability studies

Colon cancer cell lines were grown in arginine free media (Thermofisher, Cat # A2493901, US Bio Cat# D9803-07B) supplemented with 10% dialyzed FBS (ThermoFisher, Cat# 26400044) and with arginine supplementation (Sigma Aldrich Cat #A8094). Lysine (Sigma Aldrich Cat #A8094) and bicarbonate were supplemented to DMEM reference levels (**Supplemental table 1**). Cells were plated into 96 well plates (3000 cells per well) and cell viability was assessed with a luminescence-based assay (CellTiter Glo, Promega, Cat # G7572) at 48 hours on a SpectraMax M3 plate reader (Molecular Devices). For nucleotide rescue experiments, nucleobases were supplemented at concentrations up to 10x reported physiologic concentrations (*55*) (Sigma Aldrich Cat #A2786, #C3506, #G11950, #T0895). All cell lines were periodically assessed for mycoplasma contamination by PCR for genomic DNA.

#### Cancer evolution experiments

Cell lines were grown under periods of intermittent arginine deprivation (12.5 uM) using arginine-free media (ThermoFisher, Cat # A2493901) supplemented with dialyzed FBS and arginine to desired concentrations (Gibco# 26400044). Starvation cycles consisted of 4 days starvation followed by rescue with standard DMEM and dialyzed FBS. Cell lines were starved for a total of 8 cycles. In parallel, cell lines were maintained under standard tissue culture conditions and passaged to control for underlying genetic drift. At the end of the starvation cycles, DNA was extracted from both the starved and unstarved cancer cell lines (Qiagen DNeasy Blood and Tissue Kit, Cat#69506) with RNAse A treatment (Qiagen Cat# 19101) and sent for whole genome sequencing at the New York Genome Center.

#### Arginine deprivation experiments

DMEM media with varying levels of arginine were prepared as described above. Cells were plated to approximately 20% confluence in standard DMEM media supplemented with 10% v/v fetal bovine serum. At approximately 40% confluence, cells were washed three times with equal volume PBS and media was replaced with either control media (DMEM with standard amino acid concentrations and 10% v/v dialyzed FBS) or treatment media (DMEM with 12.5 uM arginine with 10% v/v dialyzed FBS). Sample collection methods for respective experiments, i.e., western blots, northern blots, etc., are described in the respective sections. Unless specified otherwise, cell samples were collected at 24 hours after initiating starvation for downstream experiments.

### Knockdown experiments

For knockdown of RARS, either SMARTPool (Horizon Discovery cat# L-009820-02) siRNA or control non-targeting siRNA (Horizon Discovery cat#D-001810-10) were utilized with Lipofectamine RNAiMAX transfection reagent (Invitrogen). Transfections were carried out 20 nM siRNA following the manufacturer’s instructions. In brief, siRNAs were diluted in Opti-MEM I (Invitrogen) and mixed with Lipofectamine RNAiMAX for 20 minutes. The mixture was subsequently added to adherent cells, rinsed with PBS (Corning), and incubated for 6 hrs before changing back to fresh complete media. Transfections were incubated for 4 days before RNA and protein collection.

#### Western blots

Protein lysates were extracted with ice-cold RIPA buffer supplemented with protease and phosphatase inhibitors (Roche). Thirty μg of protein lysates were separated using SDS–polyacrylamide gel electrophoresis and transferred to a PVDF membrane (Immobilion-P, Millipore, IPVH00010). After blocking the membranes in 5% BSA in TBST (1× TBS (Cell Signalling); 0.1% Tween20 (Sigma)), the membranes were incubated overnight at 4 °C with primary antibodies either rabbit anti-ASNS antibody (Proteintech, 14681-1-AP) diluted 1:1,000 in 5% BSA (Sigma), mouse anti-β-Actin antibody (Millipore Sigma A5441) diluted 1:5,000 in 5% BSA (Sigma), and rabbit anti-RARS (Proteintech, 27344-1-AP) diluted 1:1,000 in 5% BSA (Sigma).

Primary antibodies were incubated in 5% BSA in TBST overnight at 4 °C. After washing the blots 3 times for 15 min each in TBST, the membranes were incubated with HRP-conjugated goat anti-rabbit IgG (H+L) or HRP-conjugated goat anti-mouse IgG (H+L) secondary antibody (Invitrogen). Finally, the membranes were incubated with ECL western blot substrate (Thermo Scientific) for 1 min. X-Ray films (Fujifilm) were exposed to the western blot membranes and developed with a film processor (SRX-101A, Konica Minolta) and exposure.

### RNA isolation and purification

RNA was extracted from cells using TRIzol (Invitrogen) and isopropanol precipitation according to manufacturer’s instructions. After precipitation, the RNA pellet was washed twice with ice-cold freshly prepared 75% EtOH and then subsequently air-dried and resuspended in TE buffer.

#### Northern blots

Purified RNA was run on 10% TBE-Urea gels at 200V for 1 hour and transferred to a Hybond-N+ membrane (GE) at 150A for 1 hour. RNA was crosslinked to the membrane using UV radiation at 240 mJ/cm^2^. Membranes were blocked with Oligo Hybridization Buffer (Ambion) for 1 hour at 42 °C. Northern probes were labeled with ^32^P ATP with T4 PNK (NEB) and purified with a G25 column (GE healthcare). Probes were hybridized in Oligo Hybridization Buffer overnight at 42 °C. Membranes were washed with 2X SSC 0.1% SDS buffer and 1X SSC 0.1% SDS prior to exposing film. Films were developed with exposure times adjusted based on the probe signal strength. Probe sequences were as follows: tRNA^Arg^_UCG_: 5’-GCCTTATCCATTAGGCCACGT-3’, tRNA^Arg^_UCU_: 5’-ATCCATTGCGCCACAGAGCC-3’, tRNA^Arg^_ACG_: 5’-CCGTAGTCAGACGCGTTA-3’, tRNA^Arg^_CCG_: 5’-CCGGAATCAGACGCCTTAT-3’, tRNA^His^_GUG_: 5’-AACGCAGAGTACTAACCACTATACG-3’, tRNA^Tyr^_GUA_: 5’-ACAGTCCTCCGCTCTACCAGCTGA-3’, tRNA^Leu^_UAG_: 5’-CTCCGAAGAGACTGGAGCCTAAA -3’, and U6: 5’-CACGAATTTGCGTGTCATCCTT-3’. Multiple probes were tried for tRNA^Arg^_CCU_, however none yielded any detectable signal despite large yields of total RNA and strong signals for the other arginine tRNA. There is currently no clearly identified gene for tRNA^Arg^_GCG_ and therefore northern blots were not attempted for this tRNA(*56, 57*). Membranes were stripped by washing with 0.1% SDS in boiling water and allowing to cool to room temperature. Subsequent probes were applied starting with re-blocking with Oligo Hybridization Buffer and repeating all downstream steps with new freshly-labeled probes. Band intensity quantification was perfumed using ImageJ with the signal in each lane being normalized to U6.

#### PDX propagation

Patient derived xenografts were propagated as previously described(*27*). In brief, within 2 hours of surgical resection, CRC tumor tissue that was not needed for diagnosis was implanted subcutaneously into NSG mice at the MSKCC Antitumor Assessment Core facility.

When the tumor reached the pre-determined end-point of 1,000 mm, the tumor was excised and transferred to the Rockefeller University. Xenograft tumor pieces of 20–30 mm^3^ were re-implanted. When the subcutaneous tumor reached 1,000 mm^3^, the tumor was excised. The rest of the tumor was chopped finely with a scalpel and placed in a 50 ml conical tube with a solution of Dulbecco’s Modified Eagle Medium (Gibco) supplemented with 10% v/v fetal bovine serum (Corning), L-glutamine (2 mM; Gibco), penicillin-streptomycin (100 U/ml; Gibco), Amphotericin (1 μg/ml; Lonza), sodium pyruvate (1 mM; Gibco) and Collagenase, Type IV (200 U/ml; Worthington) and placed in a 37°C shaker at 220 rpm for 30 min. After centrifugation and removal of supernatant, the sample was subjected to ACK lysis buffer (Lonza) for 3 min at room temperature to remove red blood cells. After centrifugation and removal of ACK lysis buffer, the sample was subjected to a density gradient with Optiprep (1114542, Axis-Shield) to remove dead cells. The sample was washed in media and subjected to a 100-μm cell strainer and followed by a 70-μm cell strainer. Mouse cells were removed from the single-cell suspension via magnetic-associated cell sorting using the Mouse Cell Depletion Kit ((130-104-694, Miltenyi), resulting in a single-cell suspension of predominantly CRC cells of human origin.

### PDX metabolite profiling analysis

Metabolite profiling results from PDX samples were acquired from previously published data from our group(*27*) and processed identically to the publication.

### Metabolite extraction and profiling

Metabolite extraction and subsequent Liquid-Chromatography coupled to High-Resolution Mass Spectrometry (LC-HRMS) for polar metabolites of cells was carried out using a Q Exactive Plus and in collaboration with the Proteomics Resource Center at Rockefeller University. For all metabolite profiling, cells were washed with ice cold 0.9% NaCl and harvested in ice cold 80:20 LC-MS methanol:water (*v/v*). Samples were vortexed vigorously and centrifuged at 20,000 *g* at maximum speed at 4°C for 10 min. Supernatant was transferred to new tubes. Samples were then dried to completion using a nitrogen dryer.

Dried polar samples were resuspended in 60µL pre-chilled 50%(v/v) acetonitrile/water resuspension solvent, vortexed for 10 seconds, centrifuged for 10 minutes at 4°C and 13,200 resolution per minute, then 14µL from each sample was transferred to create a pooled sample. This pooled sample was further diluted with 1:3 and 1:10 dilution factors and employed as biological quality control. Samples were analyzed in randomized order and at 5µL injection volume via LC-MS system.

Polar metabolites were separated on a SeQuant® ZIC®-pHILIC 5µm polymer (150 mm × 2.1 mm) column (EMD Millipore) connected to a Thermo Vanquish ultrahigh-pressure liquid chromatography coupled to a Q Exactive Plus Hybrid Quadrupole-Orbitrap mass spectrometer (Thermo Fisher Scientific) with a heated electrospray ionization source. Chromatographic separation was achieved by mixing mobile phase A consisted of 20mM ammonium carbonate with 0.1%(v/v) ammonium hydroxide (adjusted to pH 9.3 with formic acid) and mobile phase B of acetonitrile in the following gradients: 90% - 40% B (0-22 min), held at 40% B (22-24 min), 40%-90% B (24-24.1 min), and reequilibrated at 90% B (24.1-30 min) at a flow rate of 0.15mL/min. Mass spectrometric data were acquired in polarity switching mode for both MS1 (full MS) and MS2 (data-dependent acquisition) with the following parameters: spray voltage, 3.0kV; capillary temperature, 275°C; source temperature, 250°C; sheath gas flow, 40 a.u.; auxiliary gas flow, 15 a.u. The full MS scans were acquired with 70,000 resolution, 1×106 ACG target, 80ms max injection time and a scan range of 55-825 m/z. The data-dependent MS/MS scans were acquired at a resolution of 17,500, 1×105 ACG target, 50ms max injection time, 1.6Da isolation width, stepwise normalized collision energy of 20, 30, and 40 units, with 8s dynamic exclusion and a loop count of 2.

Relative quantification of polar metabolites and its isotopologue was performed in Skyline Daily (v.21.2.1.403)(*58*) with the maximum mass error and retention time tolerance set to 2ppm and 12s respectively, referencing in-house retention time for polar metabolite standards. Correction to heavy isotopes were done using theoretical abundance obtained with enviPat R package (*59*). The isotopologue probabilities were defined and pruned to their absolute abundance. The peak table generated from manual peak picking from Skyline platform were cleaned up to report only prevalent polarity for each metabolite.

### Ribosome profiling

Cell lysis, ribosome footprint purification, and downstream library construction were performed according to previously published protocols 24 hours after starvation(*22*) with minor modifications. Namely, for harvesting the cells, the dishes were flash frozen on liquid nitrogen after washing them with prechilled 1x PBS, and subsequently the frozen cells were scraped off in cold lysis buffer on ice. Additionally, due to the phasing out of the legacy Ribo-Zero Gold rRNA Removal Kit from Illumina, we used the RiboCop rRNA Depletion Kit for Human/Mouse/Rat (HMR) V2 (Legoxen, cat. # 144.24) to deplete ribosomal RNA, following the manufacturer’s instructions to retrieve small RNAs after the rRNA depletion using alcohol precipitation instead of the kit purification steps. In parallel, whole RNA were also isolated for downstream RNA sequencing and normalization for translational efficiency analysis using the TruSeq RNA Library Prep Kit v2 (Illumina).

### Bioinformatics processing of ribosomal profiling experiments

For bioinformatics processing, cutadapt was used to remove the linker sequence AGATCGGAAGAGCAC(*60*) and the FastX-Toolkit (RRID: SCR_005534) used to split reads by their barcodes(*61*). Prior to further downstream analysis, reads aligning to ribosomal RNA sequences were discarded using STAR(*62*) and remaining reads were aligned to the transcriptome (GRCh38.p13). UMI-tools was used to extract unique molecular identifiers introduced during the sequencing steps and deduplicate the reads(*63*). The riboWaltz library was utilized to quantify ribosome A-site localization(*64*). Ribosomal protected footprint (RPF) counts were normalized using median of ratios normalization prior to calculating differences in A-site codon abundances in each group.

For loess regression and quantification of stalling bias, the Ribolog package was used to quantify stalling coefficients under fed and starvation conditions separately using the default spanning parameter (*23*). In this framework, local regression is utilized to smooth out peaks introduced by stalling events and measure the degree to which ribosomal stalling occurs at any given position along a transcript. Larger coefficients correspond to more stalling, with values centered around 1. All transcripts with less than three aligned RPFs were removed prior to downstream analysis. To compare amino acid or codon-specific stalling coefficients, each gene was assigned its maximum stalling coefficient for each amino acid or codon to estimate the greatest degree of stalling at a specific codon or amino acid-type for each gene. Bias coefficient ratios were calculated by taking the ratio between coefficients in starvation and fed conditions. In order to calculate amino acid identities in close proximity to maximal sites of stalling, for each gene positions were ranked by CELP bias coefficient to identify regions where maximal stalling was predicted to occur and then amino acid frequencies within the first three codons upstream and downstream of the maximum stalled site were counted. In order to restrict observations to the open reading frame, only positions that were at least ten codons downstream of the start codon and five codons upstream of the stop codon were considered for analysis. Observations were normalized for gene-specific codon content and scaled to window width to account for codon composition of each individual gene. Translational efficiency analysis was performed using CELP-corrected RPF counts.

For allele-specific ribosome profiling, mutations from whole exome sequencing were used to identify genes with heterozygous single nucleotide variants (SNV). Corresponding variant sequences were created from the wild type sequence using a custom Python transcript and subsequently added to the original reference transcriptome fasta file. Alignment, ribosome A-site localization, and quantification of stalling coefficients were performed following the same procedure as above. Comparison of ribosome stalling around single nucleotide variants were then calculated by comparing stalling coefficients upstream and downstream of the SNV and statistical analysis performed only on genes containing that were heterozygous for a SNV.

### Proteomics

The RKO cell line was selected for proteomics experiments due to its high frequency of arginine-codon switching mutations. At 24 hours after starvation at 12.5 uM arginine, cells were washed with PBS and treated with 0.25% trypsin to detach cells from the plate. Suspensions were immediately placed on ice with full media and centrifuged at 4°C, then resuspended in PBS twice to wash out media and FBS components. Following washing steps, cell pellets were resuspended in lysis buffer consisting of 0.02 M Tris-HCl (pH 7.4), 0.1 M KCl, 0.001 M EDTA (pH 8), and 0.5 M NP-40. One tablet of 1x cOmplete protease inhibitor (Roche) was added to 10 mL of fresh lysis buffer. Samples were incubated on ice and vortexed every five seconds for a total of 15 minutes, then sonicated on ice with a 4x five-second pulse at 40% amplitude with a 30 second break between samples. Following sonication, samples were transferred to clean tubes and spun down again at max speed at 4°C for 10 minutes. The resulting supernatant was transferred to an additional set of clean tubes for further proteomics analysis. Prior to any downstream processing, protein was quantified using the Pierce BCA Protein Assay Kit (cat# 23225). Twenty-five micrograms of protein from each sample was run at 200 V for 50 minutes in a 4-12% Bis-Tris gel with MOPS buffer. The gel was stained and visualized with SimplyBlue Safestain (Thermofisher cat# LC6060) following manufacturer’s instructions to ensure distinct protein bands and rule out obvious residual contamination from trace bovine serum albumin before proceeding to downstream quantification.

Further sample processing was then performed in collaboration with the Proteomics Resource Center at Rockefeller University: 50 µg of protein from each sample was reduced and alkylated using dithiothreitol and iodoacetamide. Proteins were precipitated using chloroform/water/methanol extraction and pellets were digested with Endopeptidase LysC (Wako Chemicals) and sequencing grade modified trypsin (Promega). Peptides were labeled with TMTpro isobaric tags (Thermo Scientific), pooled, purified using an Oasis HLB cartridge (Waters), and fractionated using a high-pH fractionation spin column kit (Pierce). Fractionated peptides were separated across a 2.5-hour linear gradient on a 250mm*75µm Easyspray column using a Dionex 3000 HPLC system operating at 300nL/min and analyzed by a Q-Exactive HF mass spectrometer (Thermo Scientific) operating in positive data-dependent acquisition mode. Raw data was queried against the human proteome (downloaded from uniprot.org on 2/12/2019) at 1% FDR using MaxQuant v. 1.6.1.0. Data was searched using standard settings. Further statistical analysis was performed within the Perseus framework using version 1.6.5.0. Protein-group intensities were log2-transformed and normalized by subtraction of the median. Statistical significance was tested for using FDR-corrected (permutation-based with 250 randomizations) t-test (q=0.05).

Pathway enrichment for proteomics changes was performed with iPAGE(*9*). For downstream quantification of mutational events in the proteome, proteins that were significantly increased or decreased in either fed or starvation conditions were selected with FDR < 0.05. Mutational status in each gene set was then cross-referenced to previously collected whole exome sequencing data to determine mutational status of differentially abundant proteins. Mutational changes were normalized to total number of new mutations per cell relative to the unselected cell lines.

### Whole exome sequencing and analysis

DNA was extracted using the DNeasy Blood and Tissue Kit (QIAGEN) following manufacturer’s instructions. Prior to sequencing, DNA was subjected to quality control with Picogreen and Fragment Analyzer to determine DNA integrity. Cancer samples were then sent for sequencing at the New York Genome Center. Whole exome sequencing (WES) libraries were prepared using the Agilent SureSelect XT library preparation kit in accordance with the manufacturer’s instructions. Briefly, DNA was sheared using a Covaris LE220. DNA fragments were end-repaired, adenylated, ligated to SureSelect oligo adapters, and amplified by PCR. Exome capture was performed using the Agilent SureSelect XT Human All Exome v6 (60Mb) capture probe set and captured exome libraries were ligated to Agilent Sequencing adapters during target selection and enriched by PCR. Final libraries were quantified using the Qubit Fluorometer (Life Technologies) or Spectromax M2 (Molecular Devices) and Fragment Analyzer (Advanced Analytical) or Agilent 2100 BioAnalyzer, and were sequenced on an Illumina NovaSeq 6000 sequencer run across two lanes of an S4-300 cycle flow cell.

For cancer cell lines, base calling and filtering were performed using current Illumina software; sequences were aligned to NCBI genome build 37 using Burrows-Wheeler Aligner (*65*). Picard was used to mark duplicate reads (Picard v1.83; http://picard.sourceforge.net); local realignment around insertions and deletions and base quality scores were recalibrated using GATK (Genome Analysis Toolkit v3.5, PMID: 21478889). Variants were called using GATK HaplotypeCaller, which generates a single-sample GVCF. To improve variant call accuracy, multiple single-sample GVCF files were jointly genotyped using GATK GenotypeGVCFs, which generates a multi-sample VCF. Variant Quality Score Recalibration (VQSR) was performed on the multi-sample VCF, which adds quality metrics to each variant that can be used in downstream variant filtering.

For mouse PDX exome sequencing, base calling and filtering were performed using current Illumina software. Mouse reads were then detected and removed from the FASTQ files by aligning the data to a combined reference of mouse (GRCm38) and human (NCBI genome build 37). All read pairs with both reads mapping to mouse or one read mapping to mouse and the other unmapped were excluded from subsequent processing and analyses steps. The samples were then processed through NYGC’s somatic pre-processing and variant calling pipelines. The samples were aligned to build 37 using Burrows-Wheeler Aligner (BWA-MEM v0.7.15) (*65*); NYGC’s ShortAlignmentMarking (v2.1) is used to mark short reads as unaligned (https://github.com/nygenome/nygc-short-alignment-marking). GATK (v4.1.0) FixMateInformation is run to verify and fix mate-pair information, followed by Novosort (v1.03.01) markDuplicates to merge individual lane BAM files into a single BAM file per sample. Duplicates are then sorted and marked, and GATK’s base quality score recalibration (BQSR) is performed. The final result of the pre-processing pipeline is a coordinate sorted BAM file for each sample. Variants were called using GATK HaplotypeCaller, which generates a single-sample GVCF. To improve variant call accuracy, the GVCF files were genotyped using GATK GenotypeGVCFs, and Variant Quality Score Recalibration (VQSR) was performed which adds quality metrics to each variant that was used in downstream variant filtering.

Variants were annotated using ANNOVAR(*66*). Any mutation appearing in the majority (at least 2 out of 3) control cell lines were considered parental mutations. Variant calls that did not fall into the former category were used for further analysis. In situations where mutational events were predicted to affect multiple transcripts and result in more than one possible amino acid/codon switching event, all mutational events were first counted and then normalized to the total number of transcripts affected from the single mutation. For cell line analyses, mutational counts were averaged across triplicates in each cell line and amino acid / codon switching events normalized to the total number of new single nucleotide variations in each specific sample. For PDX experiments, we made the following adjustment: since the parental tumors were generally not propagated across multiple mice in contrast to the highly-liver metastatic derivative, unique mutations were filtered out using the matched tumors as a reference. We then calculated the rate of events in shared mutations and compared this to the frequency in the mutations unique to only liver-metastatic tumors.

### Software libraries used

Experiment and model schematics were created with BioRender.com Specialized software libraries used for gene expression, ribosome profiling, and WES analyses are cited in their respective methods sections. For statistical analysis and plotting not specifically referenced, we used the following Python libraries: NumPy (1.19.2), pandas (1.13), SciPy (1.5.2), bioinfokit (2.0.3) and Seaborn(0.11.0) as well as the following R libraries: corrplot (0.92). Graphpad Prism (9.1.2) was also used to assist with statistical analysis and figure creation.

### Statistical analysis

Statistical analyses (t-tests, Mann-Whitney tests) were carried out using Prism 9 or SciPy (*67*) and were two-tailed tests unless otherwise specified in the text. Bioinformatics analyses for gene expression and ribosome profiling were carried out using specialized software packages as described under their corresponding sections. Throughout all figures, *p < 0.05, **p < 0.01, and ***p < 0.001, ****p < 0.0001. Significance was concluded at p < 0.05.

## Acknowledgments

We are grateful to members of our laboratory for comments on previous versions of the manuscript as well as Kivanç Birsoy at Rockefeller University and Balaji Santhanam at Columbia University for their suggestions. We thank Rockefeller University resource centers for assistance: Vaughn Francis and other veterinary staff of the Comparative Bioscience Center for animal husbandry and care, Henrik Molina and staff at the Proteomics Resource Center for assistance with proteomics quantification and metabolite profiling. The authors also acknowledge collaborators at the New York Genome Center for their support in processing whole exome sequencing samples. The authors would also like to acknowledge Andrew Azzam for his assistance in processing proteomics samples as a medical rotation student. The results published here are in part based upon data generated by the TCGA Research Network: https://www.cancer.gov/tcga.

## Funding

Damon Runyon Physician Scientist Training Award PST-31-20 (DJH)

National Institutes of Health Grant 5R01CA257153 (ST)

Black Family Metastasis Research Center (SFT)

Pershing Square Innovation Fund (SFT) Reem-Kayden Award (SFT)

Author contributions

Conceptualization: DJH, ST, SFT

Methodology: DJH, ST, SFT

Validation: DJH, NM, JG

Formal analysis: DJH, SH, HA

Investigation: DJH, JG, NY, AP, ML, SH, HA

Supervision: DJH, SFT, ST

Writing – original draft: DJH, SFT

Writing – review & editing: DJH, ST, SFT

### Data availability

RNA sequencing and ribosome profiling results will be made publicly available on GEO (#GSE213472) upon publication. A reviewer token has been generated and will be shared with the editor/reviewers upon request. Proteomics data will be uploaded to PRIDE for public sharing and will be available to editorial reviewers upon request. A reviewer link for the whole exome sequencing data is available at the NCBI SRA at: https://dataview.ncbi.nlm.nih.gov/object/PRJNA880559?reviewer=3rh6p2dpd1f12qcj58cm2t7bar and will be made publicly available upon publication. All custom code will be made available by the corresponding author on reasonable request.

