## supplemental_data for "Arginine limitation causes a directed DNA sequence evolution response in colorectal cancer cells"

Dennis J. Hsu, Jenny Gao, Norihiro Yamaguchi, Alexandra Pinzaru, Nandan Mandayam, Maria Liberti, Søren Heissel, Hanan Alwaseem, Saeed Tavazoie*, and Sohail F. Tavazoie*

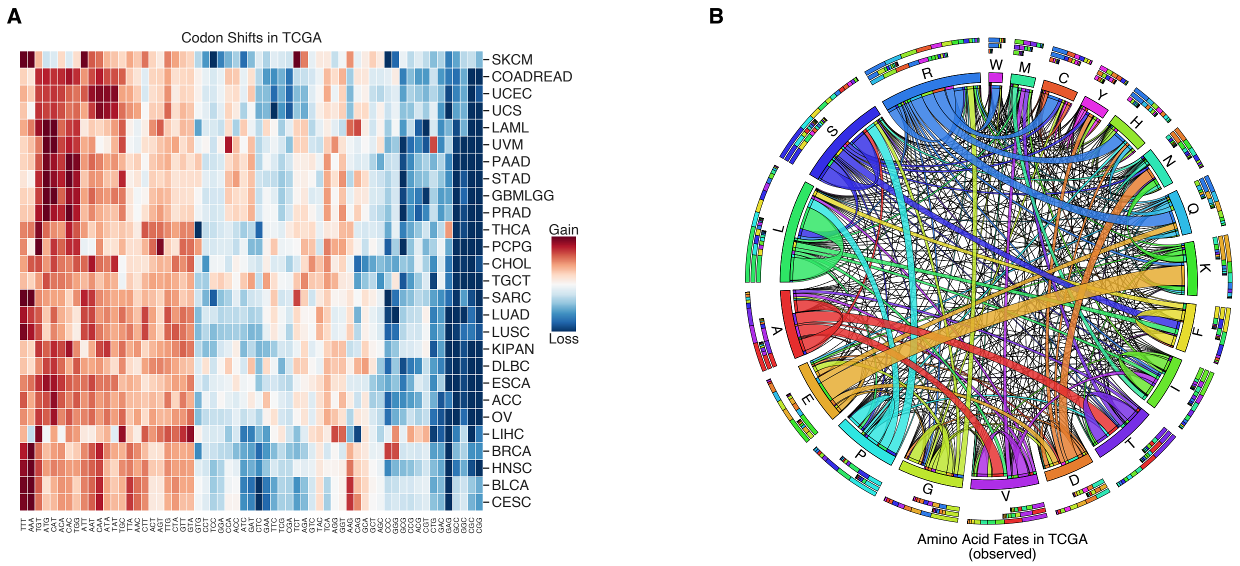


**Fig. S1. Codon changes and amino acid switching events observed across all cancers. (A)** Heatmap depicting codons gained (red) and lost (blue) across the TCGA. Gains and losses are normalized to the total number of missense and silent mutation events for each cancer type **(B)** Quantitative chord diagram depicting observed mutations pooled across all TCGA. Ribbons which directly touch a column segment indicate loss of that specific amino acid codon during a mutational event and gain of the corresponding amino acid codon in which the ribbon terminates. Ribbons that begin and end at the same amino acid represent synonymous mutations.

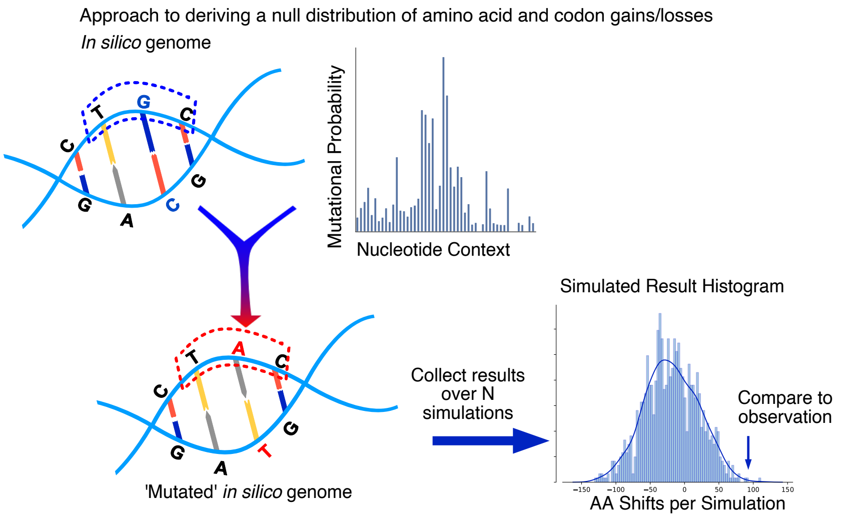


**Fig. S2. Idealized schematic of the computational approach used to study codon-switching events.** Codon and amino acid switching events were modeled *in silico* using the reference human genome and measured mutational frequencies. A histogram of potential observations was constructed (unless otherwise stated in the manuscript, N = 1000 simulations) and then compared to observed changes in the TCGA.

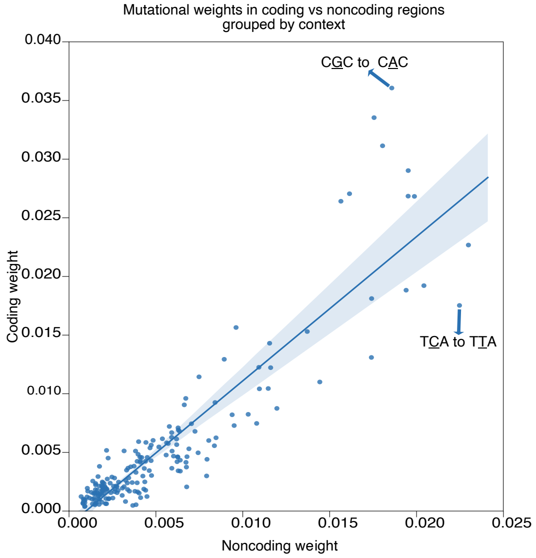


**Fig. S3**. **Mutational weights in noncoding versus coding regions of the genome show different biases**. Each point represents a specific nucleotide context and weighted frequency of change based on mutations observed in coding regions versus noncoding regions of the human genome. Two points are labeled as examples: in the first, a G to A transition is more frequent in coding regions when the nucleotide is flanked by C bases. In the second, a C to T transition, is more common in noncoding regions of the genome when the nucleotide is flanked by 5'-T and 3'-A.

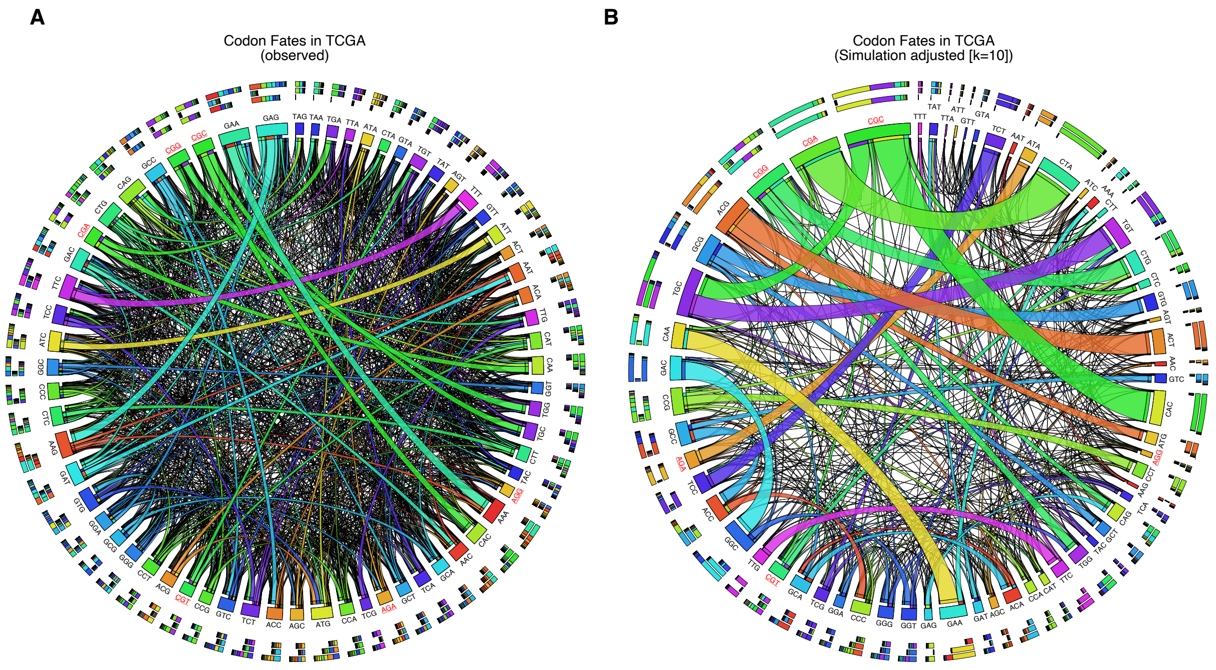


**Extended Data Fig. 4. Codon switching events with and without simulation adjustment**. **(A)** Quantitative chord diagram depicting codon shifts observed across all TCGA tumor types without scaling. Ribbon sizes are proportional to number of events. Ribbons which directly touch a column segment indicate loss of that specific amino acid during a mutational event and gain of the corresponding amino acid in which the ribbon terminates **(B)** Qualitative chord diagram showing codon shifts across all TCGA tumor types relative to neutral mutation model. (scaling factor K=10). Arginine codons are emphasized with red text.

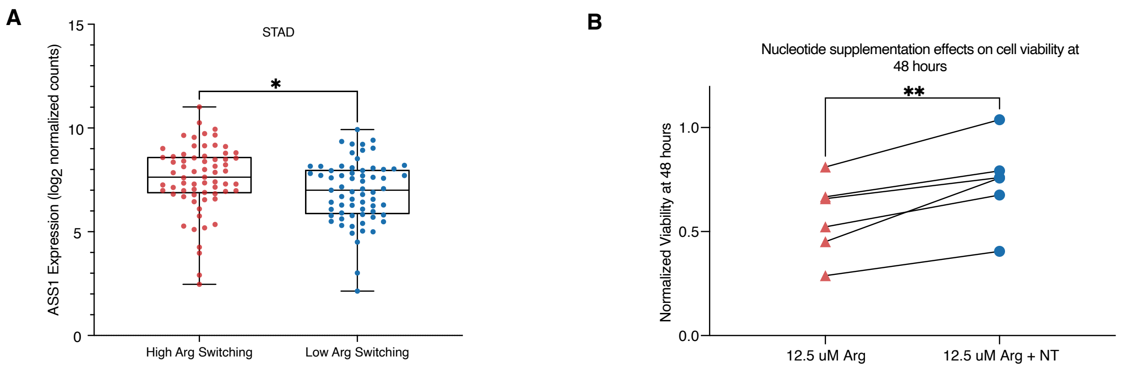


**Fig. S5. Tumors with increased arginine codon mutations are more sensitive to arginine deprivation (A)** ASS1 expression in gastric cancer samples in high-arginine codon switching versus low-arginine switching groups (DESeq2 p-adjusted *P < 0.05) **(B)** Meta-analysis of nucleotide supplementation effects on colon cancer cells following arginine deprivation, with or without nucleotide supplementation. Each individual point represents the average reading for a cell line (two-tailed paired t-test). (**P<0.01)

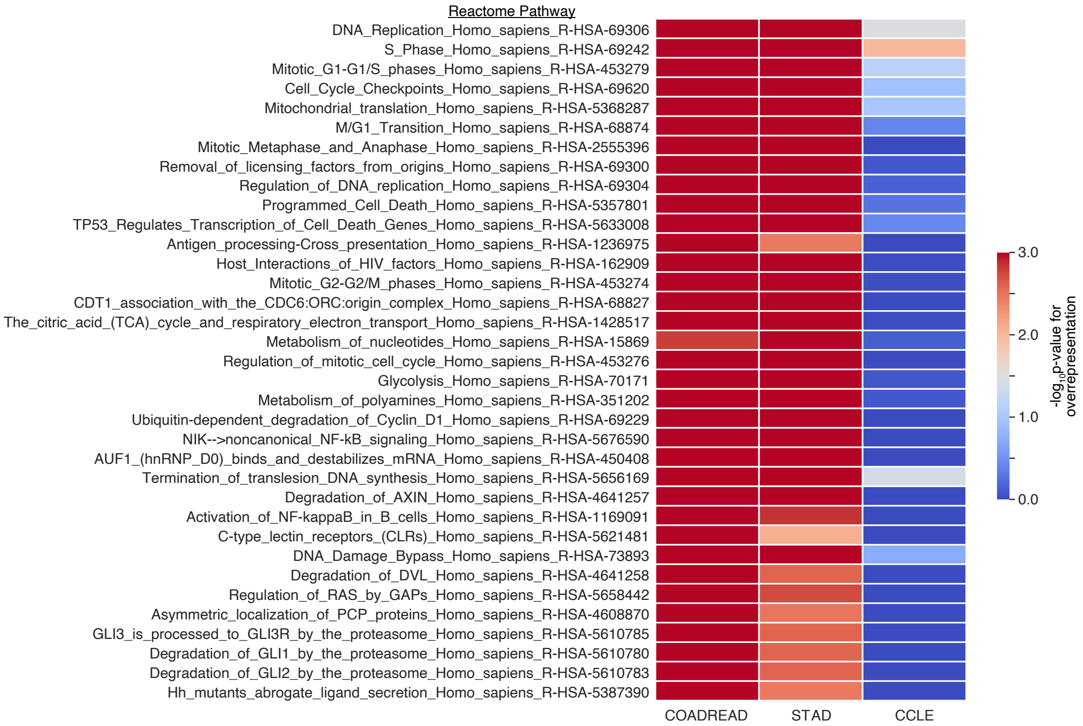


**Fig. S6. Nucleotide metabolism and bioenergetic pathways are over-represented in high arginine-codon losing tumors but not in cell lines cultured in media containing excess amino acids**. Gene expression analyses of common over or under-expressed pathways in cancer cell lines (CCLE) and colon/stomach adenocarcinoma samples from the TCGA in samples exhibiting large degrees of arginine codon-switching.

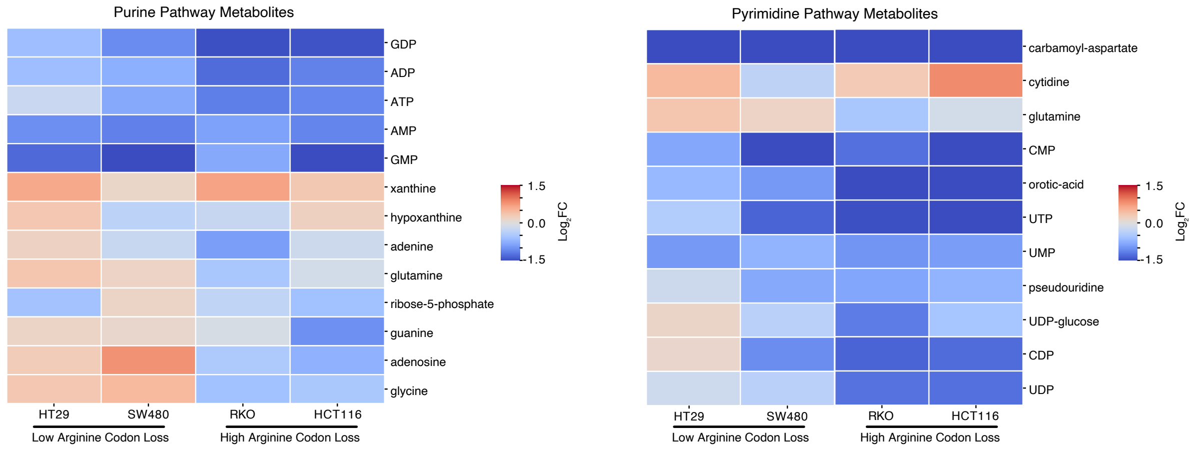


**Fig. S7. Heatmap of nucleotide metabolite changes following arginine deprivation.** Colors correspond to log2FC of annotated metabolite comparing fed versus starved cells at 24 hours after initiating starvation.

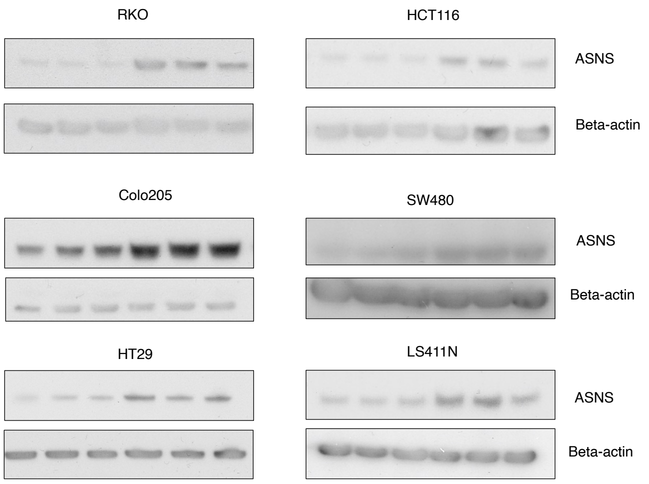


**Fig S8. Arginine deprivation results in induction of ASNS.** Western blot for ASNS following arginine deprivation in multiple colorectal cancer cell lines.

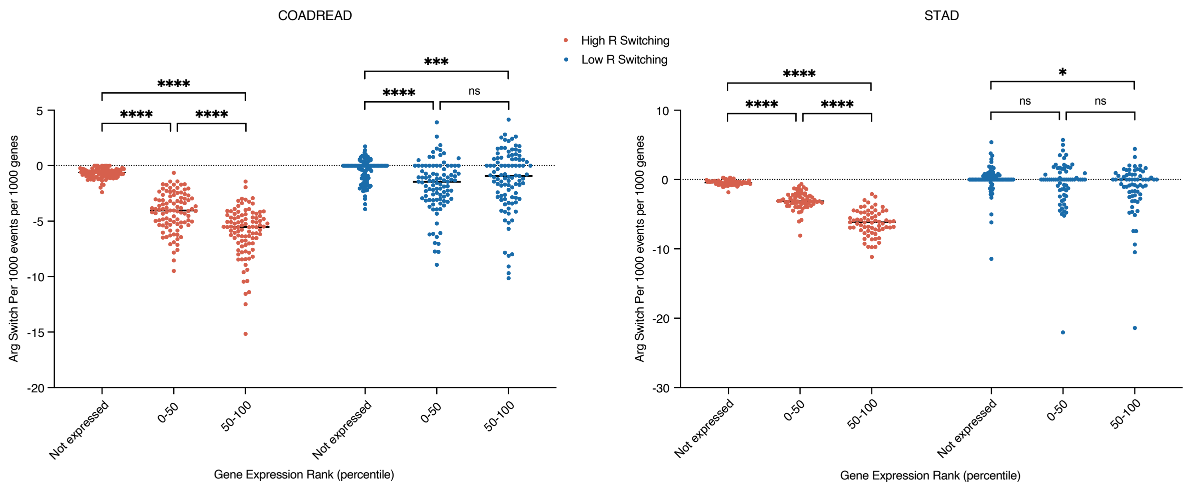


**Fig. S9. Arginine codon mutations occur more frequently in highly expressed genes.** Gene expression within individual tumors were stratified into top or bottom half expression and arginine codon mutations were counted in each group. Each dot represents the calculated value for a specific tumor in the TCGA. Statistical significance between groups was determined using a (paired two-tailed t-test). (*P<0.05, ***P<0.001, ****P<0.0001)

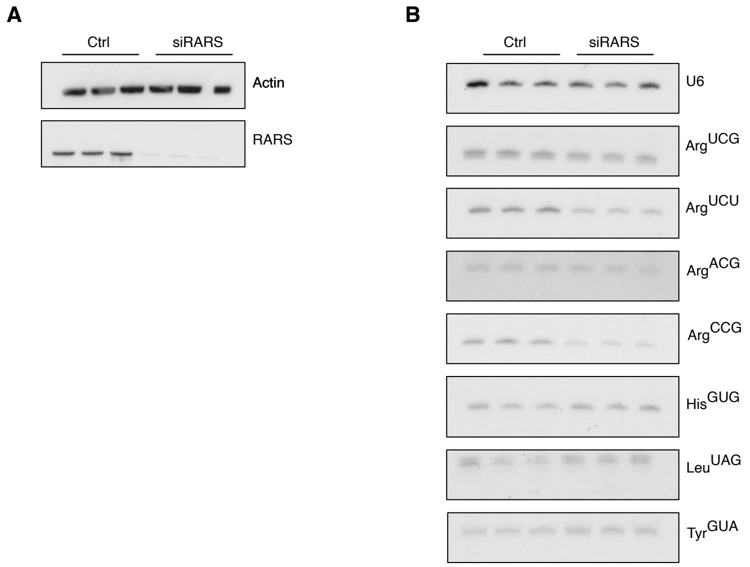


**Fig. S10. RARS knockdown phenocopies the effects of arginine deprivation on arginine tRNA abundance (A)**Western blot for RARS in cells treated with control siRNA or RARS siRNA **(B)** Transfer RNA abundance at 4 days post transfection as measured via northern blot.

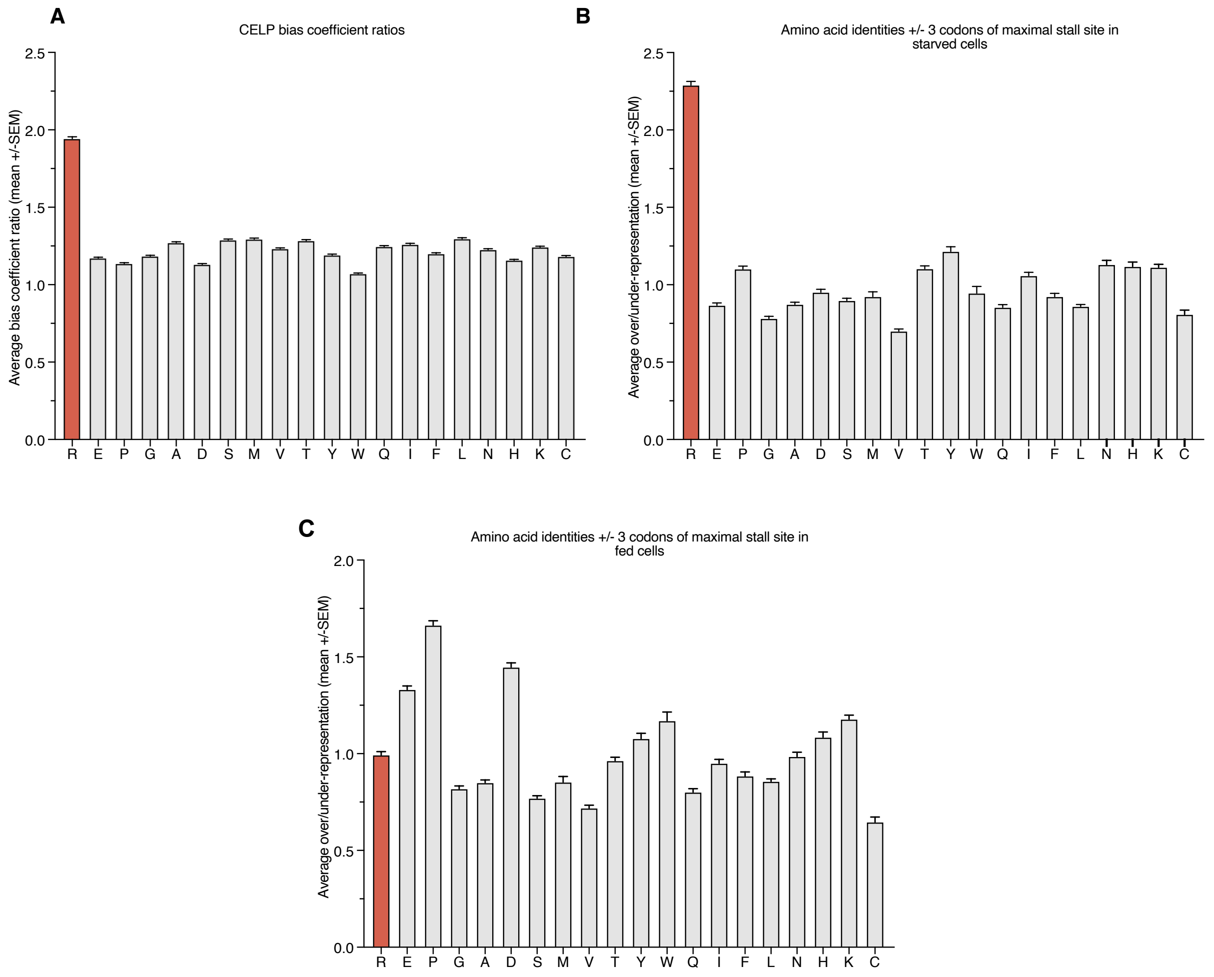


**Fig. S11. Arginine deprivation results in significantly increased stalling around arginine codons under starvation conditions**. **(A)** Average bias coefficient ratios (starved / fed coefficients, mean +/- SEM) per amino acid per gene between starved and fed conditions. Coefficients greater than 1.0 indicate more stalling under starvation conditions **(B)** Average amino acid representation near maximal stalling sites in gene transcripts under arginine deprivation conditions. Amino acid identities 3 codons upstream and downstream of the maximal stall site were calculated and normalized to gene-specific codon/amino acid usage **(C)** Average amino acid representation at sites of maximal stalling under fed conditions.

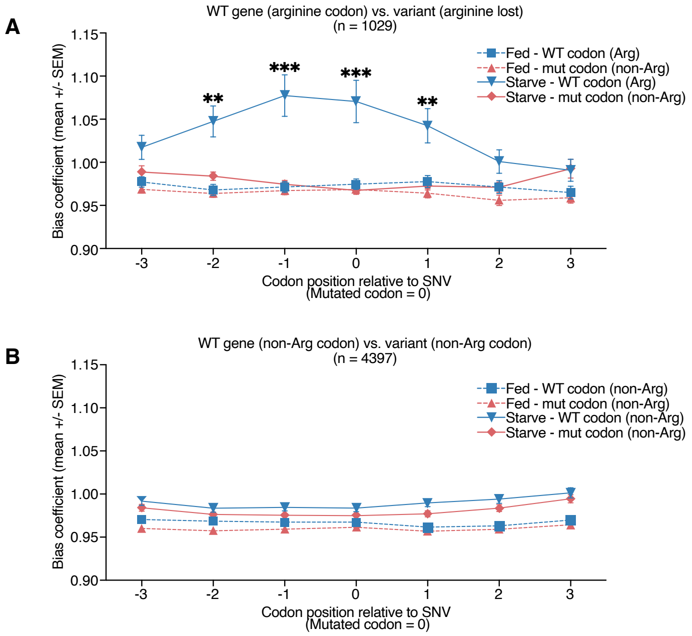


**Fig. S12. Single nucleotide variants that result in arginine-codon switching directly impact ribosome localization during arginine deprivation (A)** Bias coefficients near wild type versus variant codons when a wild type arginine codon is lost via a single nucleotide variation (SNV) **(B)** Bias coefficients upstream and downstream of all other wild type and variant codons when the substitution does not involve arginine codons. *P* values calculated using unpaired two-sided t-tests between starved conditions at each position and then adjusted using Benjamini-Hochberg with $q \leq 0.05$. (*$q<0.05$, **$q<0.01$, ***$q<0.001$.)

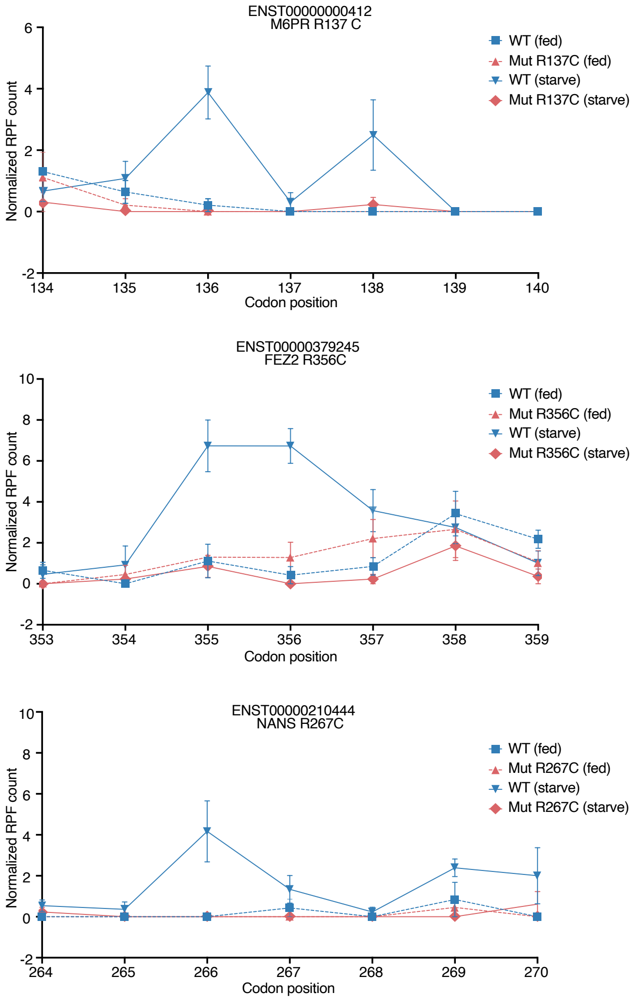


**Fig. S13. Arginine-losing SNVs impact ribosome localization**. Examples of ribosome footprint localization in heterozygous genes that lost arginine codons due to single nucleotide changes.

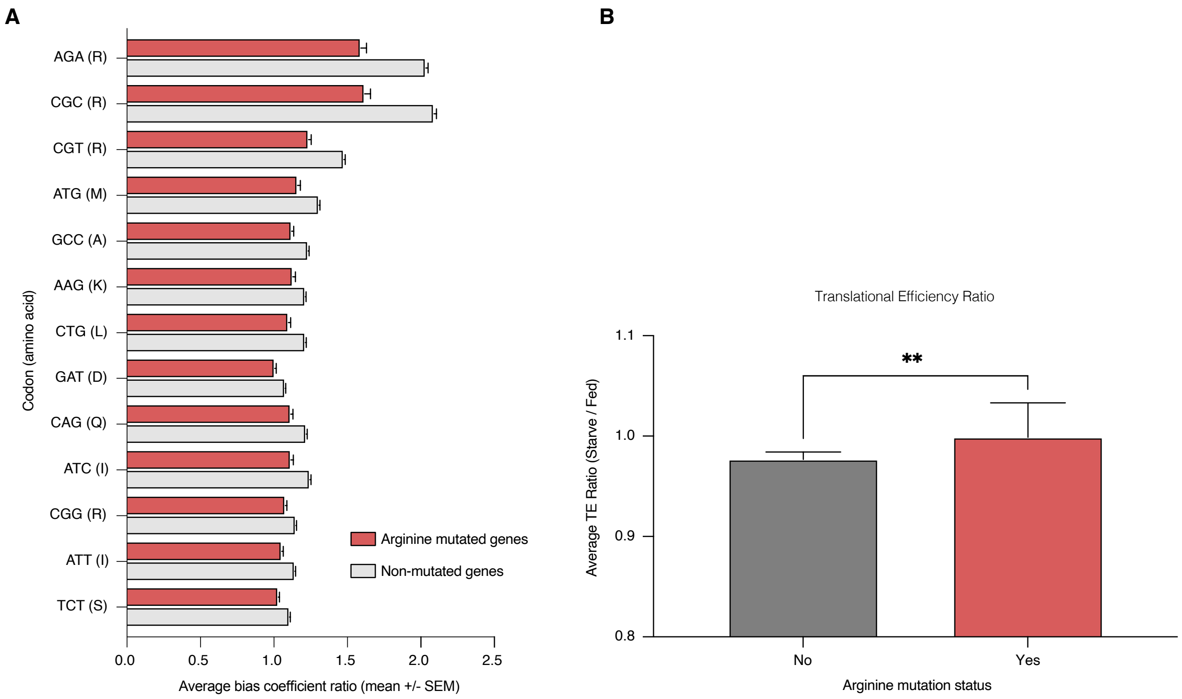


**Fig. S14. Arginine mutated genes show less evidence of ribosome stalling under arginine deprivation**. **(A)** Bias coefficient ratios (starved / fed coefficients, mean +/- SEM) in genes with or without arginine codon mutations. For ease of plotting, only statistically significant comparisons with FDR $\leq$ 0.05 (Benjamini Hochberg) are shown and all depicted codons in this diagram were considered statistically significant. Larger values imply increased stalling under starvation. **(B)** Translational efficiency ratios (starved / fed,) with CELP-bias corrected counts in genes with or without arginine codon mutations shown as median +/- 95% CI (two-tailed Mann Whitney test). (**P<0.01)

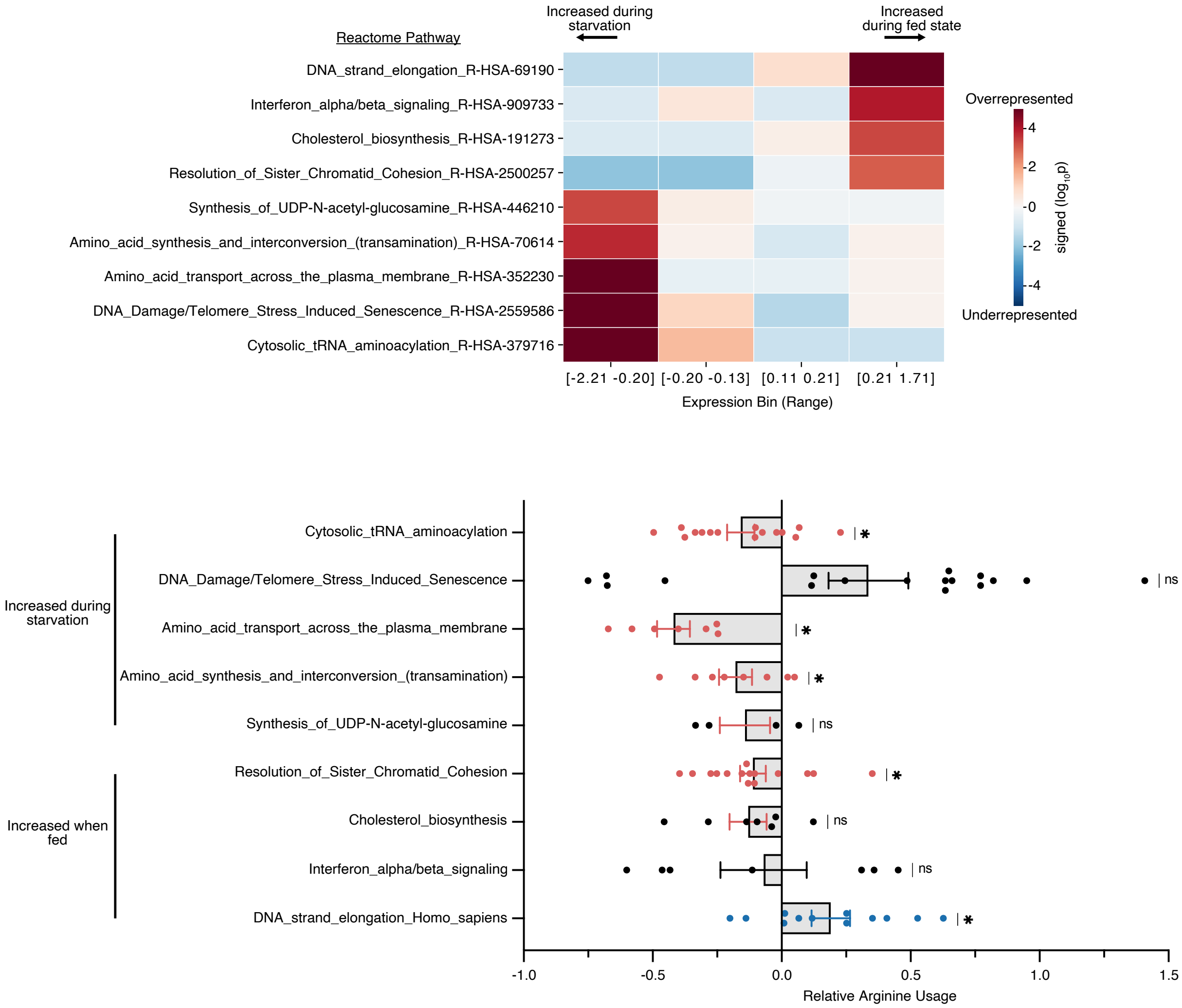


**Fig. S15. Arginine deprivation results in increased expression of DNA damage-related and amino acid synthesis/transport pathways. (A)** Protein lysate was collected from fed and starved cells, and sent for proteomics analysis. Mutual information analysis was performed to identify over-represented pathways. To simplify the diagram, only pathways with p-values < 0.001 in the most extreme bins (highly abundant under starvation or during fed states) are shown. **(B)** Relative arginine usage for proteins upregulated in fed and in starved states. Each point represents relative arginine usage (relative to all other proteins in the human genome) for a specific gene in the pathway. Bar height represents mean in each group with error bars indicating SEM. (One sample Wilcoxon test with $\mu0 = 0$, i.e. no preference for or against arginine usage). (*$P<0.05$)

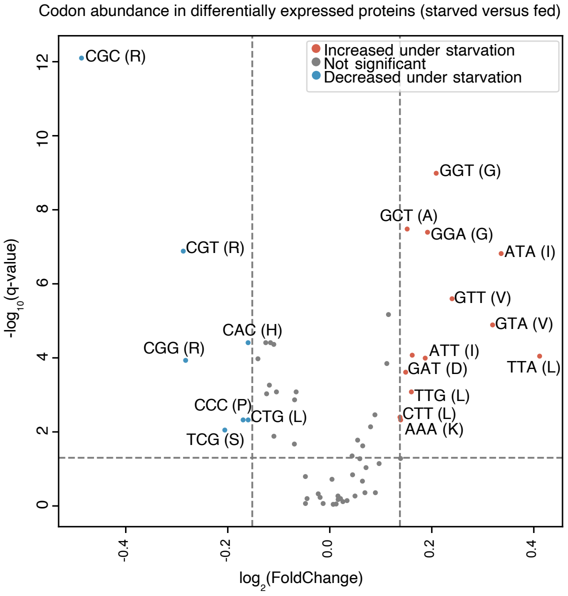


**Fig. S16. Arginine codon usage in the tumor proteome following starvation**. Codon usage of genes expressed under starved conditions relative to fed states.

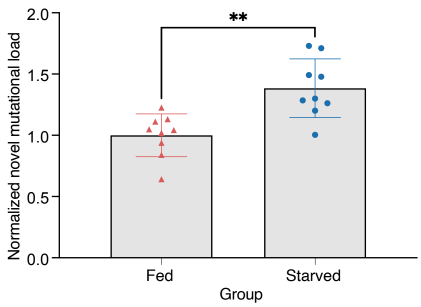


**Fig. S17. Arginine deprivation increases rates of mutation.** Novel mutational loads in individually starved cell lines. Each point corresponds to a cell line replicate. Mutational loads are normalized to new mutations acquired in the fed group (Two-tailed Mann-Whitney test). (**P<0.01)

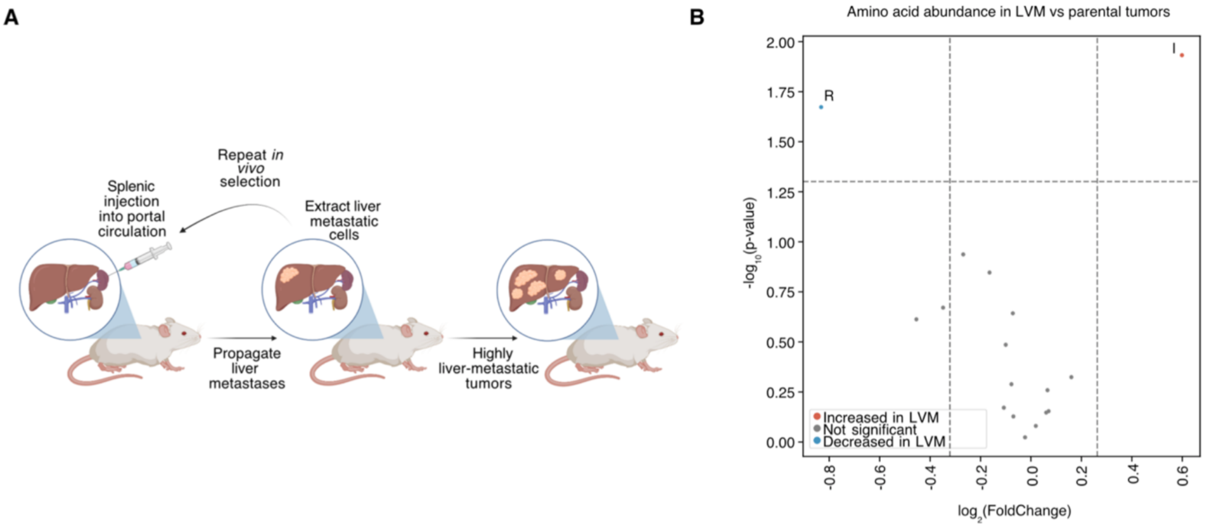


**Fig. S18 . Free arginine is low in highly liver metastatic PDX tumors. (A)** Schematic of how *in vivo* highly-liver metastatic tumors were derived, and **(B)** LC-MS based metabolic profiling on highly liver metastatic (LVM) tumors versus parental tumors. Only amino acids are depicted on the volcano plot.

| Component | DMEM (mM) | Serum (mM) | Ratio (Media:Serum) |
|---|---|---|---|
| L-Arginine * HCl | 0.399 | 0.074 | 5.39 |
| L-cystine*2HCl | 0.2 | ** | N/A |
| L-glutamine | 3.996 | 0.536 | 7.46 |
| Glycine | 0.4 | 0.275 | 1.45 |
| L-Histidine * HCl * H2O | 0.2 | 0.071 | 2.82 |
| L-Isoleucine | 0.8 | 0.08 | 10.01 |
| L-Lysine*HCl | 0.799 | 0.12 | 6.39 |
| L-Methionine | 0.201 | 0.053 | 3.79 |
| L-Phenylalanine | 0.4 | 0.112 | 3.57 |
| L-Serine | 0.4 | 0.151 | 2.65 |
| L-Threonine | 0.798 | 0.106 | 7.52 |
| L-Tryptophan | 0.078 | 0.045 | 1.74 |
| L-Tyrosine * 2Na * 2H2O | 0.397 | 0.077 | 5.16 |
| L-Valine | 0.802 | 0.196 | 4.09 |

**Reference serum values inferred from:** Le, A., Ng, A., Kwan, T., Cusmano-Ozog, K. and Cowan, T.M., 2014. A rapid, sensitive method for quantitative analysis of underivatized amino acids by liquid chromatography–tandem mass spectrometry (LC–MS/MS). Journal of Chromatography B, 944, pp.166-174.

**Table S1. Amino Acid concentrations in standard media versus reported values measured in human serum.**

| Cell Line | Net arginine codon changes | Assigned group |
|---|---|---|
| HCT116 | -279 | High loss |
| LS411N | -326 | High loss |
| RKO | -170 | High loss |
| Colo205 | -20 | Low loss |
| SW480 | -38 | Low loss |
| HT29 | -31 | Low loss |

**Table S2. Panel of colorectal cancer cell lines.**
